# HERACLES for fast and accurate likelihood-based reconstruction of massive lineages

**DOI:** 10.64898/2026.11.06.764882

**Authors:** Sitara Persad, Quaid Morris

## Abstract

CRISPR-based lineage recording technologies can trace the ancestry of hundreds of thousands of cells, yet reconstructing accurate phylogenies at this scale remains computationally challenging. Likelihood-based methods employ explicit evolutionary models of CRISPR editing that improve tree accuracy but require computationally expensive searches over tree topologies, whereas greedy distance-based alternatives scale efficiently but their accuracy requires distances that faithfully represent the true phylogeny. Here, we introduce HERACLES, which optimizes cell coordinates in hyperbolic space under a quartet-based composite likelihood of CRISPR editing. By recovering local branching structures directly from embedded distances, HERACLES guides continuous optimization with an explicit evolutionary model without repeatedly constructing and evaluating complete phylogenies. Across simulated benchmarks, HERACLES improves reconstruction accuracy over existing methods and scales to lineages containing 150,000 cells in about four hours on a single GPU. On experimental KP-tracer and PEtracer datasets, HERACLES reconstructs phylogenies consistent with independent anatomical annotations and temporal lineage markers. Its flexible likelihood framework also accommodates different recording mechanisms, and incorporating sequential editing improves reconstruction of simulated DNA Typewriter lineages. Together, these results establish continuous optimization of local evolutionary relationships as a scalable and flexible approach to probabilistic lineage reconstruction.

**Code availability:** HERACLES is available at https://github.com/morrislab/HERACLES.

## 1 Introduction

Most phylogenetic inference algorithms are designed for trees with hundreds to thousands of leaves. Recently developed lineage tracing methods create new challenges because they can generate data for hundreds of thousands of cells and they have unique, technology-dependent mutation processes. For example, CRISPR-based recorders introduce irreversible, heritable mutations into engineered genomic targets allowing cell ancestry and molecular phenotypes to be profiled with the same assay[1–3]. Faithful recovery of the corresponding phylogeny would precisely link phenotype and shared ancestry at single cell resolution, which has clear, transformative applications in biology [4–8]. Current phylogeny reconstruction methods are ill-suited to this major computational challenge.

Existing methods face a trade-off between accuracy and computational tractability. The most accurate models attempt exhaustive [9, 10] or stochastic search [11–13] over tree space for high scoring trees under either parsimony or likelihood criteria. These approaches scale poorly because the search space grows super-exponentially with the number of cells. Greedy optimization algorithms – neighbour joining (NJ), UPGMA or Cassiopeia-Greedy [9, 14, 15] – are computationally efficient but inaccurate. While NJ will recover the correct phylogeny when given a matrix of the pairwise tree distances between all leaves [16], these distances are not readily apparent from raw data, and computing them accurately requires modeling the tree. Here, we show that it is possible to efficiently estimate accurate pairwise tree distances under an explicit evolutionary model without reconstructing a full phylogeny, thus allowing us to combine the accuracy of likelihood-based inference with the efficiency of NJ. Our algorithm uses leaf node embeddings, which are optimized using stochastic gradients, and leverages the close link between trees and hyperbolic space geometries.

Gradient-based optimization has increasingly been used in place of discrete combinatorial search [17–20]. Gradient-based phylogenetic inference can be achieved by representing cells as points in a continuous space and optimizing their coordinates so that the embedded distances approximate tree distances, e.g., evolutionary distances that maximize a likelihood[17–19]. VINE uses this approach for CRISPR-based lineage reconstruction, using distances between embedded cells to construct trees scored under an explicit CRISPR-based evolutionary model. However, its loss function requires reconstructing a complete tree at every optimization step, a computation that is at least quadratic in the number of cells at each iteration, severely limiting scalability even though the search itself is continuous.

One way to avoid the cost of tree construction is to evaluate evolutionary likelihoods directly from embedded distances, without reconstructing the complete tree. Wilson [21] demonstrated this approach using a composite likelihood of cell pairs in hyperbolic space. However, independently scoring pairs does not directly constrain their relationships through a shared branching structure and, as we demonstrate, leads to substantially less accurate reconstructions. A separate challenge is the numerical instability of optimizing tree embeddings in hyperbolic space, particularly at large distances from the origin [22, 23]. Here, we introduce a stable likelihood-based embedding method that constrains the tree structure to ensure accurate reconstruction with NJ while avoiding building a full tree at each gradient update.

HERACLES is a scalable likelihood-based framework that optimizes cell coordinates in hyperbolic space under an explicit, user-defined evolutionary model. Hyperbolic space is well suited to representing trees because it accommodates exponential branching and can preserve tree distances with low distortion [24, 25]. HERACLES adds two key innovations to prior embedding methods: scoring likelihoods of sampled cell quartets in place of a full tree, and directly penalizing deviations from tree distance.

HERACLES uses hyperbolic distances and Gromov products to recover the topology and branch lengths of rooted quartets without explicitly representing ancestral nodes. It evaluates the likelihood of each quartet under a model of CRISPR editing and uses the resulting gradients to optimize cell coordinates. Together with regularization, each quartet constrains several distances through a shared branching structure so that optimizing overlapping quartets encourages a globally consistent, tree-like distance matrix. The resulting distances reflect the editing process and more closely approximate a tree metric, allowing us to optimize tree distances directly and recover the complete phylogeny using a single call to NJ at convergence.

## 2 HERACLES algorithm

### 2.1 Representing cell lineages in hyperbolic space

A phylogeny can be equivalently represented by its topology with branch lengths or its induced set of pairwise tree distances; NJ applied to the latter recovers the former. The distance representation is useful because distances vary continuously and can be optimized using gradient descent. However, not all distance matrices correspond to a tree; pairwise distances correspond to a phylogeny if and only if they satisfy the *four-point condition* [26]. For any quartet *Q* = {*a, b, c, d*}, order the three sums {*d*_*ab*_ + *d*_*cd*_, *d*_*ac*_ + *d*_*bd*_, *d*_*ad*_ + *d*_*bc*_} as *σ*_(1)_(*Q*) ≥*σ*_(2)_(*Q*) ≥*σ*_(3)_(*Q*). The four-point condition requires the two largest of these sums to be equal, i.e. the *four-point defect*, 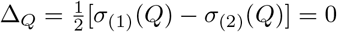.

Hyperbolic space is a natural choice for embedding trees. It readily accommodates hierarchical branching because volume in hyperbolic space grows exponentially with radius. The distances between any set of points – e.g., those representing leaf embeddings – in hyperbolic space, ℍ^*D*^, satisfy a relaxed version of the four-point condition: the maximal four point defect is at most *δ*_*H*_ [27], where *δ*_*H*_ depends only on the curvature of the space. Thus, trees can be embedded in hyperbolic space with arbitrarily low distortion, under appropriate scaling [24] and NJ is guaranteed to recover the correct tree when the input distances are sufficiently close to the true tree distance[16]. These properties have motivated hyperbolic embeddings for hierarchical representation learning [25], including the analysis of branching cell-state trajectories [28], and for phylogenetic inference [17–19, 21].

HERACLES exploits an additional property of hyperbolic spaces that permits inference of ancestral relationships directly from hyperbolic distances. For a tree with root *ρ*, the Gromov product 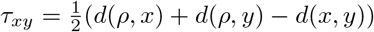 equals the depth of the most recent common ancestor of *x* and *y*. When applied to distances in hyperbolic space, the Gromov product approximates the corresponding ancestral depth in a tree representation, with additive error bounded by *O*(*δ*_*H*_ log *n*) for *n* embedded points in a *δ*_*H*_-hyperbolic space [27]. This allows us to estimate branching relationships through ancestral depths without explicitly representing ancestral nodes. The likelihood of a topology explaining the observed edits can then be evaluated on subsets of cells directly from their coordinates, without constructing a full tree.

### 2.2 Overview of HERACLES

HERACLES reconstructs rooted cell lineages from a character matrix *Y* ∈ ℤ^*n*×*S*^ containing the observed states of *S* recording sites across *n* cells (Fig. 1a). These observations are evaluated under an evolutionary model that can be adapted to different recording technologies without changing the optimization procedure. For conventional CRISPR/Cas9 recording, each entry *Y*_*is*_ = 0 represents an unedited site, *Y*_*is*_ *>* 0 identifies an edited allele, and *Y*_*is*_ = − 1 denotes a missing observation. We use the Probabilistic Mixed-type Missing model [11] to describe irreversible, heritable editing, while accounting for missing observations arising from heritable deletion or stochastic dropout and augment the character matrix with an all-unedited root, *ρ* representing the ancestral state of the lineage.

**Figure 1:**
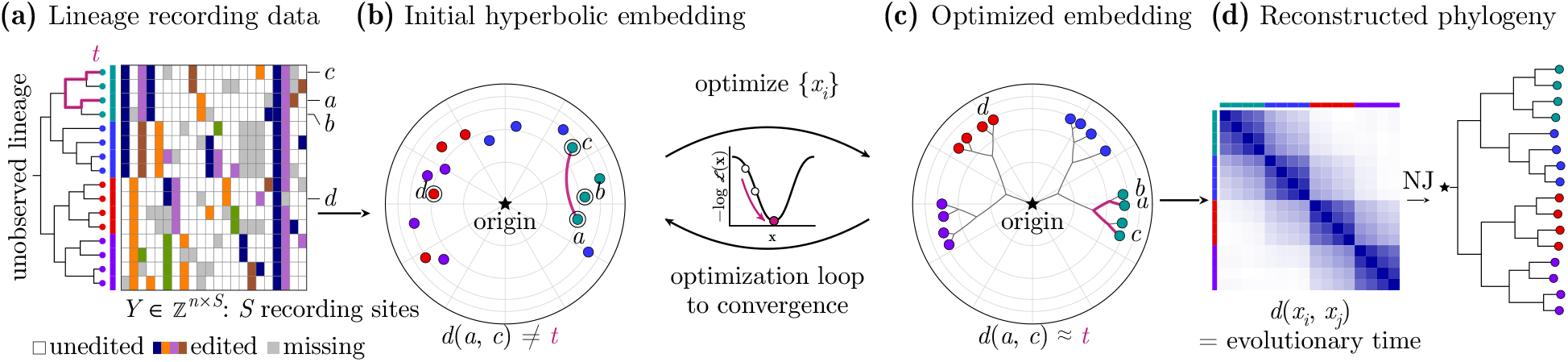
HERACLES overview. **(a)** Character matrix *Y* of observed edits and an underlying true lineage; grey marks missing observations. **(b)** Cells are initialized on a hyperboloid around the founder, which is fixed at the origin. **(c)** Cell coordinates are optimized using sampled quartet likelihoods. **(d)** Optimized distances are decoded by neighbour joining (NJ).

We represent each cell as a point in hyperbolic space and optimize its coordinates under a likelihood model of the observed edits. We use the Lorentz hyperboloid model of *D*-dimensional hyperbolic space [29], with curvature −1 and distance *d*(·, ·).

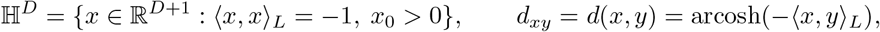

where 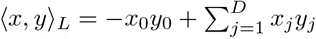. The embedding dimension *D* is selected as a function of the number of cells and is fixed during optimization (Supp. Methods A1). The unedited root cell is placed at the origin of the space and remains fixed throughout optimization (Fig. 1b).

HERACLES comprises three main steps, summarized below and elaborated in the following sections:

1. **Initialize a hyperbolic embedding**. Coordinates are initialized from the observed character matrix. Weighted-Hamming distances are used to construct a preliminary tree by neighbour joining; classical multidimensional scaling (MDS) of the resulting tree-path distances provides initial Euclidean coordinates, which are mapped onto the hyperboloid (Supp. Methods A1). A distance-preserving transformation places the founder at the origin. Note that for larger problems, where the tree building dominates the computational time, the Weighted-Hamming distances can instead be embedded directly with minimal loss in performance (Supp. Fig. S4).
2. **Optimize cell coordinates under an evolutionary model**. At each iteration, HERACLES performs three operations:
  a. *Sample informative quartets*. Select groups of four cells, preferentially within a close neighbourhood.
  b. *Evaluate quartet likelihoods*. Recover the rooted topology and branch lengths of each sampled quartet directly from the current hyperbolic distances, and calculate the probability of its observed recording states given a specified evolutionary model and the inferred evolutionary distances.
  c. *Update the embedding*. Combine the sampled quartet likelihoods with a root-to-leaf likelihood and geometric penalties that encourage tree-like distances and consistent root-to-leaf depths. The resulting objective is optimized using Riemannian stochastic gradient optimization, which updates the cell coordinates directly in hyperbolic space alongside the parameters of the evolutionary model.
3. **Reconstruct the lineage**. After optimization, recover a rooted phylogeny from the learned evolutionary distances using NJ. For massive ultrametric problems, NJ can be replaced with UPGMA.

### 2.3 Selecting informative quartets

For *n* cells, the number of possible quartets is *O*(*n*^4^); as such, HERACLES samples subsets of these quartets. Uniformly sampled quartets often contain cells whose common ancestors lie near the root, providing little information about local branching relationships. To sample branching relationships more evenly across the lineage, HERACLES uses a *neighbourhood-based sampling* strategy that preferentially selects quartets spanning neighbouring lineages (Fig. 2a). Before optimization, we use the Gromov product calculated from recorder-specific distances 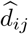 to rank potential relatives of each cell by their estimated shared-ancestor depth. These ranks are computed once at initialization and reused for quartet selection throughout optimization. For each quartet, we select an anchor cell and a neighbourhood size *k*, drawn log-uniformly up to 1,024, and then draw the remaining three cells uniformly from the anchor’s *k* nearest relatives. We then reject any quartet when the closest cell to the anchor is closer to one of the other sampled cells.

**Figure 2:**
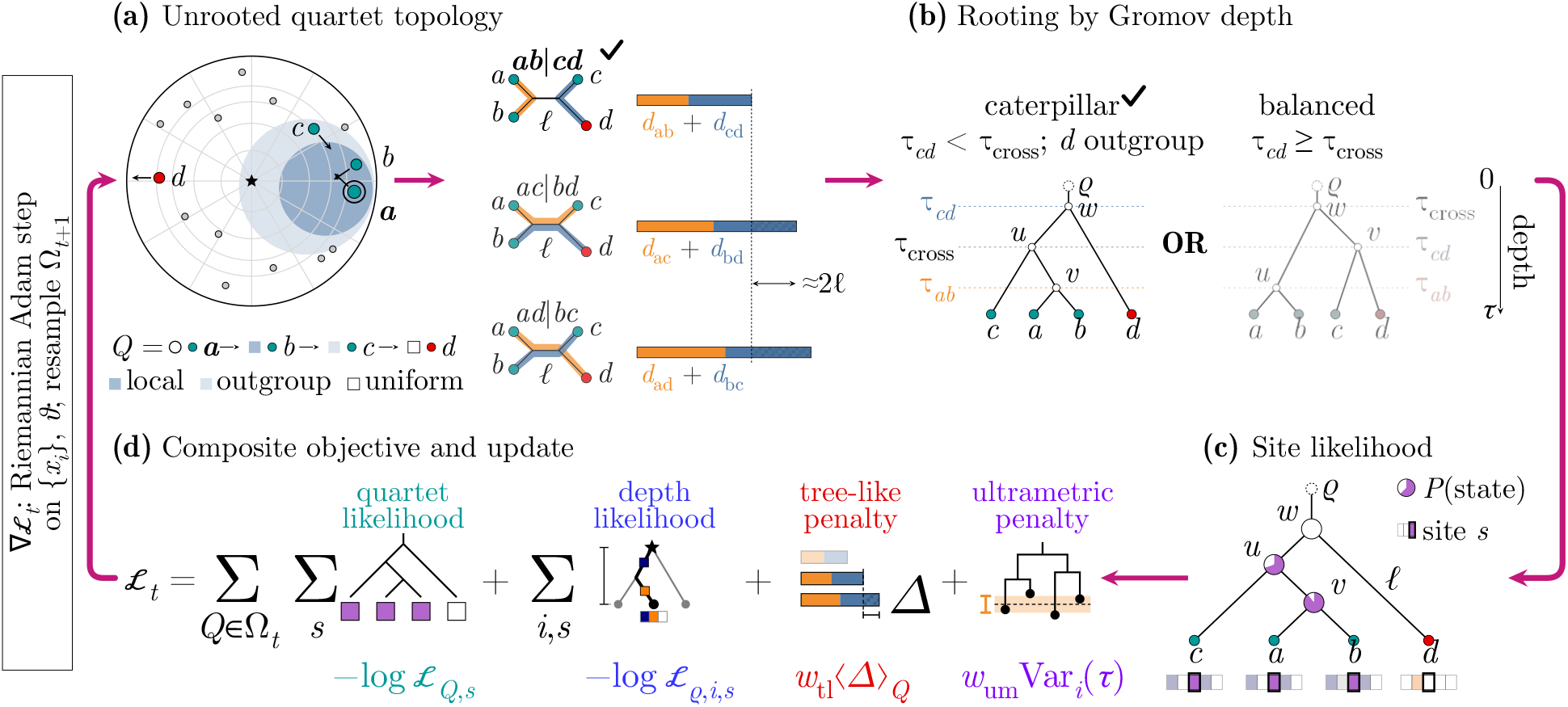
HERACLES optimization loop. **(a)** The smallest four-point distance sum selects the unrooted split *ab* |*cd*. **(b)** Gromov-product estimates of ancestral depths determine rooting and branch lengths; *τ*_*cd*_ *< τ*_cross_ selects the illustrated caterpillar with *d* as outgroup, where *τ*_cross_ is the mean cross-pair ancestral depth. **(c)** The inferred quartet topology is scored using likelihood of observed edits. **(d)** The composite loss combines negative log-likelihood contributions from quartets and root-to-leaf paths, with tree-like and ultrametric regularization. Riemannian Adam updates coordinates and model parameters.

These sampled quartets test branching relationships at multiple depths of the lineage (Supp. Fig. S9), suggesting that neighbourhood sizes span every scale from siblings to larger clades.

Then, at each iteration *t*, HERACLES samples a new set of quartets Ω_*t*_, with |Ω_*t*_| = 15*n*. In expectation, each cell is sampled in multiple quartets, each contributing gradient signal for the cell’s embedding. The number of sampled quartets grows linearly with *n*, so the cost of evaluating their likelihoods is *O*(*n*× *S*) per iteration for *S* recording sites. Further details of the sampling distributions and rejection procedure are provided in Supp. Methods A2.

For each quartet, HERACLES uses the current embedding to infer their rooted topologies and branch lengths and evaluate their likelihoods under the evolutionary model. The resulting likelihood gradients are combined with those of the root-to-leaf likelihood and geometric penalties that encourage tree-like distances and ultrametricity. HERACLES then updates the cell coordinates using Riemannian stochastic gradient optimization. Unlike the fixed distances used for quartet sampling, the hyperbolic distances used to compute quartet likelihoods change at each iteration.

### 2.4 Recovering rooted quartets from embedded distances

HERACLES infers a rooted phylogeny for each quartet from the current hyperbolic distances. We first identify the unrooted topology by comparing the three four-point distance sums and selecting the smallest. For example, if *σ*_(3)_(*Q*) = *d*_*ab*_ + *d*_*cd*_ ≤min {*d*_*ac*_ + *d*_*bd*_, *d*_*ad*_ + *d*_*bc*_}, the inferred split is *ab*| *cd* (Fig. 2a). We then root the topology by estimating the depth of the most recent common ancestors of all pairs of cells *i, j* in the quartet using their Gromov products *τ*_*ij*_, where larger values indicate more recent shared ancestry. For the split *ab* |*cd*, we compare the within-pair depths *τ*_*ab*_ and *τ*_*cd*_ with the mean cross-pair depth, 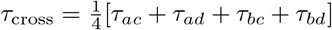. If both within-pair depths exceed *τ*_cross_ then *a, b* and *c, d* form separate clades and the quartet is balanced. A smaller within-pair depth indicates that one member of that pair branches off earlier, giving a caterpillar topology. For example, if *τ*_*cd*_ *< τ*_cross_, either *c* or *d* is the outgroup. We distinguish them by comparing their shared-ancestor depths with *a* and *b*: if *τ*_*ac*_ + *τ*_*bc*_ *> τ*_*ad*_ + *τ*_*bd*_, then *d* is the outgroup (Fig. 2b; Supp. Methods A3). Each quartet requires only six pairwise hyperbolic distances, together with distances to the founder. Since HERACLES samples *O*(*n*) quartets per iteration, recovering their topologies requires *O*(*n*) distance evaluations.

### 2.5 Scoring rooted quartets under a model of CRISPR editing

HERACLES next evaluates the likelihood of a quartet phylogeny (henceforth referred to as *quartet likelihood*) under an explicit model of CRISPR editing, marginalizing over unobserved ancestral states using Felsenstein pruning [30]. Let *T*_*Q*_ denote the inferred phylogeny for quartet *Q* and *θ* denote parameters of the mutation model. The likelihood of the observed states at site *s* is ℒ_*Q,s*_ = *P*_*θ*_(*Y*_*Q,s*_| *T*_*Q*_), where *Y*_*Q,s*_ = {*Y*_*is*_ : *i* ∈*Q*}.

For conventional CRISPR/Cas9 recording, we model irreversible editing at each target site using a continuous-time Markov chain (CTMC). Each site starts unedited and acquires allele *α* at rate *λq*_*sα*_, where *q*_*sα*_ is the probability of acquiring that allele upon editing (Fig. 2c). We fix the editing rate at *λ* = 1, since its timescale can be absorbed into the branch lengths and learn allele distributions, *q*. Conditional on no deletion, transitions along a branch of length *l* satisfy

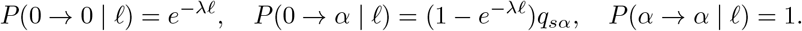

We use the Probabilistic Mixed-type Missing (PMM) model [11], which extends the star homoplasy model [10] to account for heritable deletion and stochastic dropout. Under this model, an unedited site may acquire any allele, whereas an edited site can only retain the same allele or become heritably missing. Heritable deletion occurs at rate *ε*_*H*_ along branches and is irreversible. In contrast, stochastic dropout hides the underlying state at a sampled cell with probability *ε*_*S*_ (Fig. 2c). Parameters *ε*_*H*_ and *ε*_*S*_ are estimated jointly with the embedding (Supp. Methods A4, Supp. Fig. S5).

These restrictions simplify likelihood evaluation. The root is fixed in the unedited state, and an edited ancestral state implies that this edit is present in all non-missing descendants. Similarly, heritably missing ancestral states are possible only when all descendants are missing. Thus, at most three ancestral states need to be considered during Felsenstein pruning, allowing quartet likelihoods to be evaluated quickly (Fig. 2c). Full derivations of quartet likelihoods, including the treatment of missing observations, are provided in Supp. Methods A4, Supp. Methods A5.

### 2.6 Optimizing a regularized composite likelihood

We add three regularization terms to the likelihood objective: a root-to-leaf likelihood, a penalty on depth variance, and penalty for the four-point defect. The root-to-leaf likelihood acts as a soft constraint on the depth of leaf embeddings. The depth variance term encourages ultrametricity. Overall, these regularizations encourage the embedding to be more tree-like and numerically stable.

Hyperbolic space embeddings require trading off numerical stability with tree-like-ness. The curvature of hyperbolic spaces is a tunable parameter, and increasing the magnitude of negative curvature reduces the upper bound on the four-point defect of embedded points. However, because hyperbolic coordinates grow exponentially with distance from the origin, and increasing the curvature worsens this effect, large coordinate values are difficult to represent accurately with finite-precision, leading to numerical instability [22, 23]. In HERACLES, we choose a relatively lower curvature space, and these regularizations help to shrink large coordinate values while also minimizing four-point defect, as there exist leaf configurations within the lower curvature space that can achieve much lower four-point defect than the maximum bounds. We therefore fix the curvature at 1 for good numerical stability, and penalize the average four-point defect as follows:

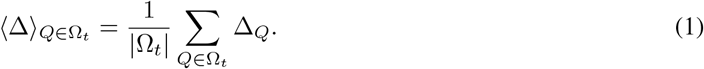

Two further loss terms act to constrain embedding depth directly. First, we include a root-to-leaf likelihood that scores each cell’s observed edits given its evolutionary distance from the founder, thereby anchoring radial distances to the editing process. At iteration *t*, let Ω_*t*_ denote the sampled quartets and *h*_*i*_ = *d*(*ρ, x*_*i*_) the root-to-leaf depth of cell *i*. For each recording site *s*, we define the root-to-leaf likelihood as ℒ_*ρ,i,s*_ = *P*_*θ*_(*Y*_*is*_ |*h*_*i*_). Combining these terms with the quartet likelihoods gives the composite loss

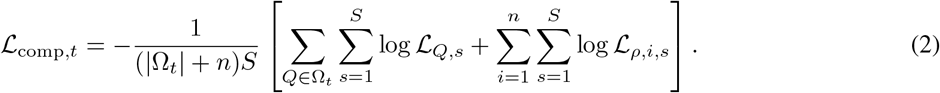

Second, we penalize variation in leaf depths using an ultrametric penalty. When editing rates are approximately constant and cells are collected contemporaneously, their root-to-leaf depths should also be similar. We define this penalty as: 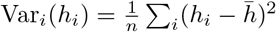, where 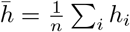.

The total loss combines these terms as follows.

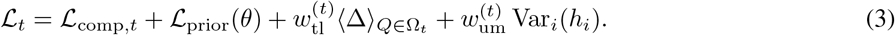

Here ℒ _prior_ is the negative log-prior on the editing and missingness parameters, including a Gamma prior on the heritable deletion rate and a Beta prior on the stochastic dropout probability (Supp. Methods A6). The coefficients 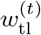 and 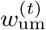 control the tree-like and ultrametric penalties, respectively. The ultrametric penalty is omitted when its assumptions do not hold.

Strong tree-like regularization early in optimization restricts exploration of alternative evolutionary relationships. We therefore optimize in two stages: first without the four-point penalty, then with its weight adjusted adaptively to meet a target value of 0.01 (Supp. Fig. S7). At each iteration, Riemannian Adam [31] updates the cell coordinates {*x*_*i*_} and learnable model parameters *θ*, after which a new quartet batch Ω_*t*+1_ is sampled (Fig. 2d).

After optimization, the embedding (Fig. 1c) is decoded into a complete phylogeny using neighbour joining (Fig. 1d), or UPGMA for very large datasets. The likelihood guides the embedding toward distances supported by the observed edits, while geometric regularization encourages those distances to describe a tree. Details of the parameter priors, adaptive penalty weights, stopping criteria and tree decoding are provided in Supp. Methods A6.

## 3 Results

### 3.1 Informative quartet sampling improves lineage reconstruction

Here, we illustrate the importance of neighbourhood quartet sampling and geometric regularization by comparing with embeddings fit using pairwise likelihoods, rooted triplets, and unregularized quartets. To illustrate the impact of incorporating the quartets, we compared embeddings optimized using all 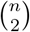 pairwise likelihoods with objectives using 30*n* sampled pairs, 20*n* triplets or 15*n* quartets per iteration. The sampled objectives evaluate the same number of cell appearances, allowing us to compare likelihood order at a matched sampling budget while avoiding the quadratic cost of evaluating all pairs. Note that the four-point defect penalty can only be assessed for quartets. To assess the value of neighbourhood-based quartet sampling and regularization, we compared uniform and neighbourhood-based quartet sampling, with and without the four-point defect penalty.

Under stochastic missingness, reconstruction accuracy improved from pairs to triplets to uniformly sampled quartets, but none outperformed neighbour joining (NJ) (Fig. 3a). In contrast, neighbourhood-based quartet sampling reduced mean Robinson–Foulds (RF) distance by 0.11 relative to uniform sampling (*P <* 10^−6^), outperforming NJ on all 25 simulated datasets. Thus, selecting informative quartets improved accuracy more than increasing likelihood order alone. Under mixed heritable and stochastic missingness, all likelihood objectives, including pairwise likelihoods, outper-formed NJ (Fig. 3b), suggesting an advantage from explicitly modeling the editing and missingness processes when missing observations can reflect inherited deletion events. Neighbourhood-based quartet sampling again produced the largest improvement, reducing mean RF distance by 0.09 relative to uniform sampling (*P* = 2 ×10^−6^).

**Figure 3:**
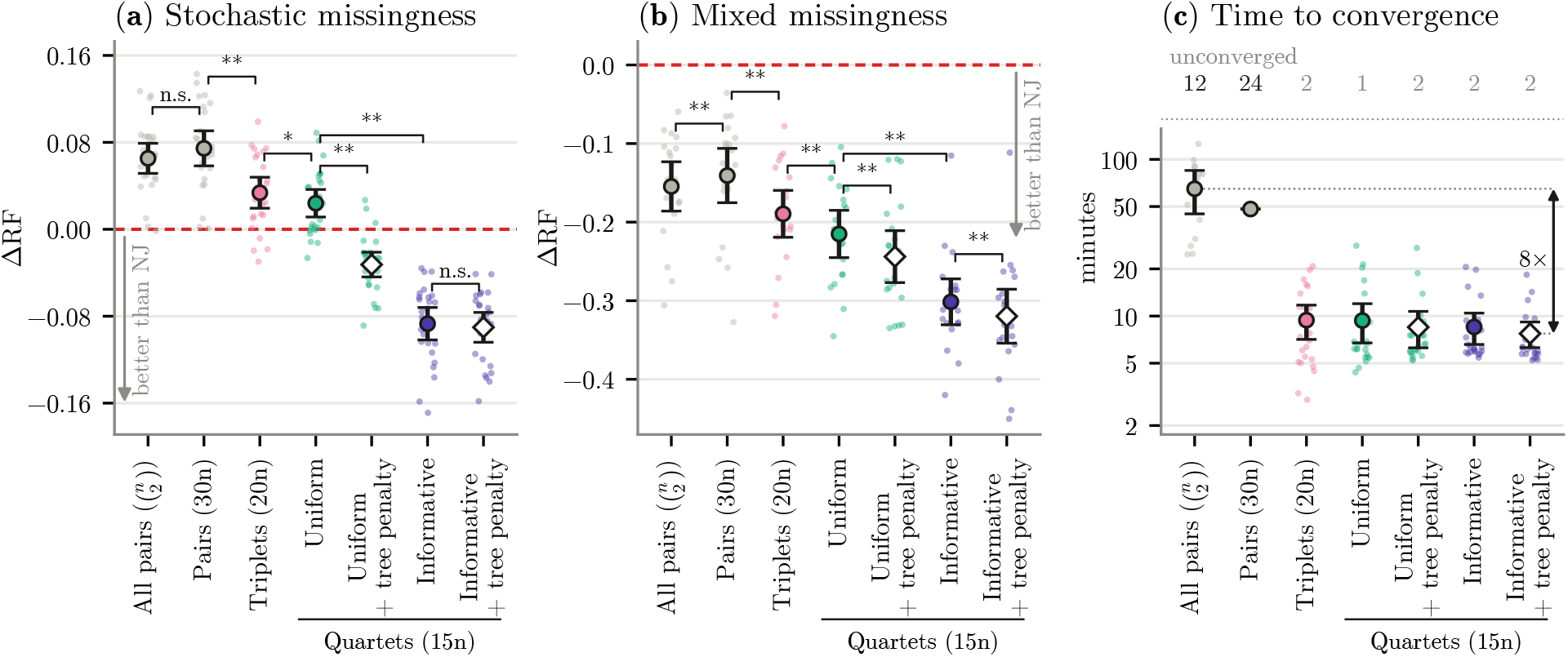
Effects of likelihood order, quartet sampling and geometric regularization on reconstruction. **(a,b)** Reconstruction accuracy relative to neighbour joining on weighted-Hamming distance (NJ) under (a) stochastic missingness and (b) mixed heritable and stochastic missingness. Negative values indicate better RF distance than NJ. Mean± 95% CI; points are datasets. Brackets: paired Wilcoxon tests (\**P <* 0.05, \*\**P <* 0.01). **(c)** Wall clock time under stochastic missingness.

The four-point penalty improved uniformly sampled quartets in both regimes (Fig. 3a,b). However, when combined with neighbourhood-based sampling, it provided little additional benefit under stochastic missingness (ΔRF = −0.003, *P* = 0.51), but significantly improved reconstruction under mixed missingness (ΔRF = −0.018, *P* = 0.003). These results suggest that enforcing tree-like distances may be particularly useful when inherited missingness complicates the inference of ancestral relationships.

Finally, we examined the computational cost of these objectives (Fig. 3c). Evaluating all pairs was substantially more expensive than scoring linearly sampled subsets. However, reducing the number of pairwise likelihood terms did not by itself yield reliable convergence: the sampled-pairs objective rarely satisfied the stopping criterion within the iteration limit. In contrast, triplet and quartet objectives typically converged much faster than the pairwise objectives. Together, these results show that accurate and efficient reconstruction depends not simply on the number of likelihood terms evaluated, but on the branching relationships they constrain. Quartet likelihoods with neighbourhood-based sampling provide the strongest reconstruction accuracy across both simulation regimes, with geometric regularization offering additional improvements in some settings (Supp. Fig. S6).

### 3.2 HERACLES enables accurate, likelihood-guided lineage reconstruction at atlas scale

We next asked whether HERACLES could retain the accuracy of likelihood-based reconstruction while scaling beyond the limits of conventional full-tree optimization. We compared HERACLES against probabilistic, parsimony-based and distance-based methods across published LAML and Startle benchmarks and independently generated lineages containing up to 150,000 cells using Cassiopeia [9] (Fig. 4). For all trees, we collapsed mutationless edges into polytomies using the Camin-Sokal procedure, and measured reconstruction accuracy using rooted Robinson–Foulds (RF) distance, with complete comparisons provided in Supplementary Tables S2–S3 and simulation details in Supp. Methods A10.

**Figure 4:**
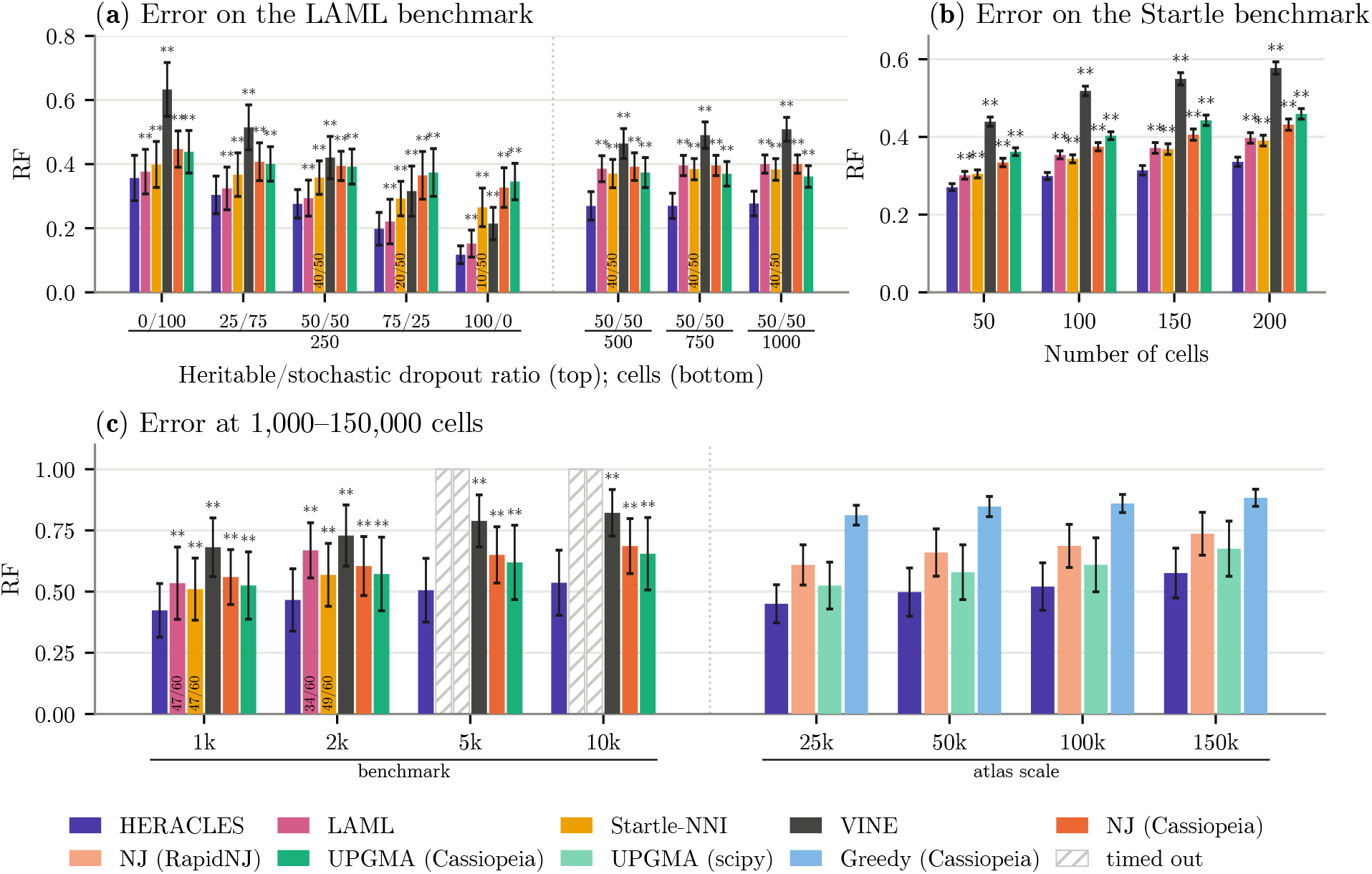
Reconstruction accuracy across simulation benchmarks and lineage scales. **(a)** LAML benchmark by missingness regime and cell count. **(b)** Startle benchmark by cell count. **(c)** Simulated lineages of 1,000–150,000 cells. Bars show mean collapsed rooted RF distance ± SD; points show individual replicates. Hatched bars indicate methods exceeding 24 hours; fractions give completed runs. Asterisks denote two-sided paired Wilcoxon tests against HERACLES (\**P <* 0.05, * * *P <* 0.01; uncorrected); no asterisk indicates *P* ≥ 0.05 where tested. Atlas-scale groups have no significance markers (*n* = 5).

HERACLES achieved the lowest mean reconstruction error across all eight conditions of the LAML benchmark (Fig. 4a). Its advantage over LAML increased with lineage size, from Δ*RF* = −0.023 at 250 cells to Δ*RF* = −0.12 at 500–1,000 cells. At these larger sizes, HERACLES improved 150/150 reconstructions, whereas LAML’s topology search returned its NJ initialization unchanged in 146/150 cases (Supp. Results B2). This suggests that the difficulty of exploring alternative tree topologies can limit the accuracy of conventional likelihood-based reconstruction even when these methods remain computationally feasible. HERACLES also achieved lower reconstruction error on the independent Startle benchmark, with mean RF distance 0.301 compared with 0.346 for Startle-NNI and 0.350 for the LAML pipeline (Fig. 4b). These improvements were observed across recording conditions (Supp. Fig. S10), indicating that HERACLES’s advantage was not restricted to a single simulation benchmark.

The scalability difference became more pronounced as lineage sizes increased. We evaluated 240 independently simulated datasets containing 1,000–10,000 cells (Fig. 4c). At 5,000 and 10,000 cells, neither LAML nor Startle-NNI completed reconstruction within the 24-hour time limit, leaving greedy distance-based methods as the principal scalable alternatives. VINE successfully scaled to these sizes, but produced significantly less accurate trees, potentially due to its likelihood model which does not separately account for heritable silencing and stochastic dropout (Supp. Table S2). HERACLES continued to reconstruct these lineages while maintaining lower error than NJ and UPGMA, reducing mean RF distance by 0.14 −0.15 and 0.10 −0.12, respectively. It outperformed both methods on 240/240 datasets. Thus, HERACLES retained the accuracy advantage of evolutionary-model-based optimization at lineage sizes where conventional full-tree likelihood methods were no longer practical.

We next tested whether this advantage persisted at atlas scale, using simulated lineages containing 25,000–150,000 cells. To accommodate these sizes, we used a low-memory configuration with 8*n* sampled quartets per iteration and an average-linkage tree read-out (Supp. Methods A8). HERACLES outperformed UPGMA, RapidNJ and Cassiopeiagreedy on 20/20 datasets, with mean RF distance of 0.576 at 150,000 cells, compared with 0.675 for UPGMA and 0.736 for RapidNJ (Fig. 4c). Importantly, its advantage over UPGMA increased from Δ*RF* = −0.075 to Δ*RF* = −0.1 across the four lineage sizes, rather than diminishing as the number of cells increased. The accuracy gains from likelihood-guided optimization therefore persisted at atlas scale.

We finally examined the computational requirements of these reconstructions (Supp. Fig. S11). GPU acceleration reduced HERACLES runtime at 10,000 cells from about 5 hours on four CPU cores to about 17 minutes on one GPU. At 100,000 cells, reconstruction required a median of 2.0 hours and 86 GB of peak host memory, compared with 0.9 hours and 81 GB for UPGMA. At 150,000 cells, HERACLES completed reconstruction in 4.1 hours using 190 GB of peak host memory, compared with 1.6 hours and 181 GB for UPGMA. Although HERACLES required more computation than distance-based methods, its peak host-memory usage was comparable to UPGMA. In both methods, peak memory was dominated by the final tree read-out, which requires storing pairwise distances, rather than by HERACLES’s hyperbolic embedding, whose coordinates occupied 2.3 GB at 150,000 cells. Thus, the principal memory bottleneck at atlas scale was tree construction rather than likelihood-guided embedding optimization.

### 3.3 HERACLES reconstructs metastatic lineages with fewer inferred migrations

We next applied HERACLES to experimental lineage-recording data from the KP-tracer mouse model of metastatic lung adenocarcinoma [7]. We analysed tumour 3724_NT_All, comprising 1,461 cells with distinct edit patterns sampled from a primary lung tumour, three liver metastases and a soft-tissue metastasis, using the same deduplicated subset analysed by LAML [11]. Although the true phylogeny is unknown, anatomical sampling locations provide information independent of the recorder edits used for reconstruction. We therefore evaluated each reconstructed tree by the minimum number of tissue transitions required to explain the observed locations, as in previous methods [11]. Because recorder edits do not resolve every branching relationship, fully bifurcating trees can contain arbitrary resolutions of polytomies that affect inferred migration counts. To avoid evaluating branching relationships unsupported by the data, we collapsed branches without support from recorder edits before calculating migration parsimony, applying the same procedure to all methods (Supp. Methods A12).

HERACLES reconstructed a tree requiring 107 migrations, compared with 120 for LAML, 123 for Startle-NNI and 136 for Cassiopeia-Hybrid (Fig. 5). The reduction was driven primarily by fewer inferred reseeding events into the primary tumour: 35 for HERACLES, compared with 54–68 for the other methods. In particular, HERACLES inferred only 30 transitions from the soft-tissue metastasis back to the primary tumour, compared with 47–57 for the other reconstructions. Conversely, HERACLES inferred 71 outward seeding events from the primary tumour, compared with 63–67 for the other methods. Thus, the lower total migration count reflects a different balance of metastatic seeding and reseeding, rather than a uniform reduction in all inferred tissue transitions.

**Figure 5:**
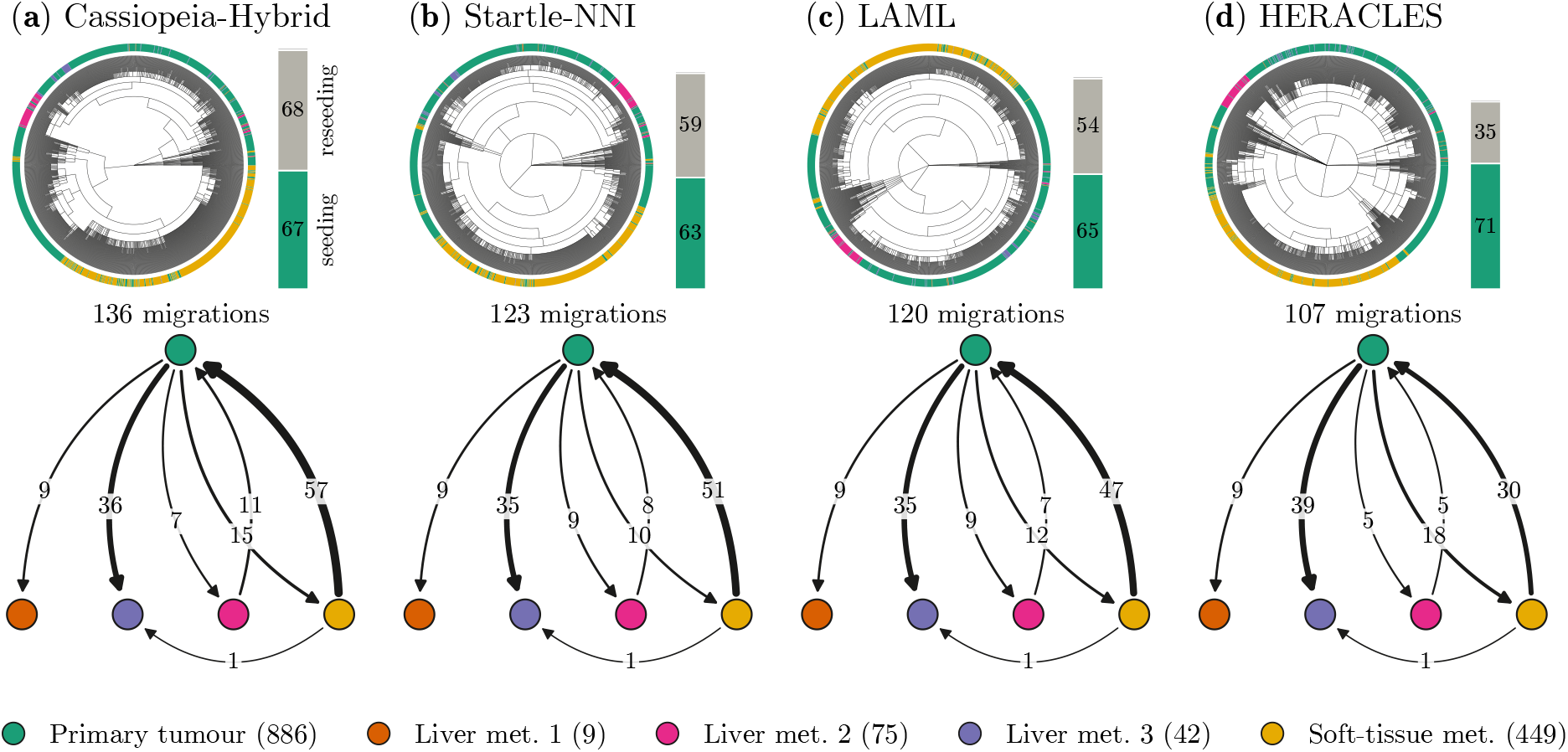
Metastatic lineage reconstruction from experimental KP-tracer data. **(a–d)** Circular phylogenies of 1,461 cells with distinct edit patterns from KP-tracer tumour 3724_NT_All, reconstructed using **(a)** Cassiopeia-Hybrid, **(b)** Startle-NNI, **(c)** LAML and **(d)** HERACLES. Outer rings indicate anatomical sampling locations, measured independently of the lineage-recording edits. Migration graphs below each phylogeny show the minimum number of anatomical-site transitions inferred by parsimony after collapsing branches unsupported by recorder edits. Arrows indicate inferred transitions between tissues, with widths and labels proportional to transition counts. Bars beside each tree summarize seeding events from the primary tumour and reseeding events into the primary tumour. Total migration counts are shown below each phylogeny.

All three independent HERACLES initializations produced trees requiring 107–108 migrations after collapsing unsupported branches (Supp. Results B3). The tree shown in Fig. 5 was selected by its fitted evolutionary likelihood, without using anatomical annotations. The comparison nevertheless depends on how unresolved branches are treated: without collapsing, LAML required slightly fewer total migrations than HERACLES (99 versus 101), although HERACLES still required fewer reseeding events (Supp. Table S5). Migration parsimony does not establish the true metastatic history, but the lower migration count indicates greater concordance between the HERACLES reconstruction and independently observed tissue locations.

### 3.4 HERACLES accommodates diverse lineage-recording models

Lineage-recording technologies differ in how edits accumulate and are inherited, requiring different evolutionary models to interpret their recordings. HERACLES allows the quartet likelihood to be adapted to these mechanisms without changing the underlying hyperbolic optimization. We evaluated this capability using two prime-editing recorders: experimental PEtracer data, in which missingness occurs at the cassette level [3], and simulated DNA Typewriter data, in which edits accumulate sequentially within each cassette [2].

We first applied HERACLES to 3,108 cells from PEtracer clone 4 (Fig. 6). PEtracer records edits at 36 sites arranged in 12 three-site cassettes, where failure to recover a cassette produces missing observations at all three sites. We therefore adapted the likelihood to model cassette-level dropout as a single event rather than three independent events. To evaluate the reconstructed lineages, we used independent lentiviral barcodes introduced on days 5 and 7, which are inherited by descendant cells but were not used for phylogenetic inference. Because barcode integration is irreversible, cells carrying the same barcode should descend from a common integration event. We therefore counted the minimum number of independent integrations needed to explain the barcode distribution on each tree [13]. If cells carrying the same barcode are separated into distinct lineages, additional integrations must be inferred. As arbitrary resolutions of polytomies can reduce the inferred integration count, we again collapsed branches unsupported by recorder edits before scoring all reconstructions, as in all other analyses.

**Figure 6:**
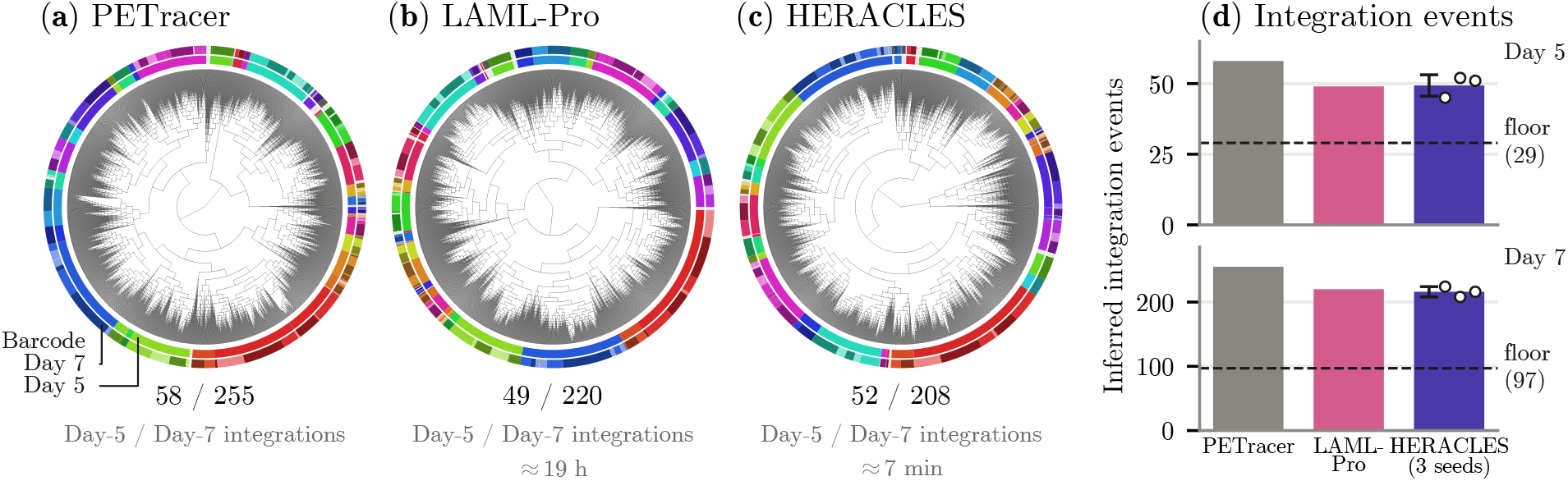
Lineage reconstruction from experimental PEtracer data. **(a–c)** Circular phylogenies of 3,108 cells from PEtracer clone 4, inferred by **(a)** PEtracer, **(b)** LAML-Pro and **(c)** HERACLES. Inner and outer rings indicate day-5 and day-7 lentiviral barcodes, respectively; grey indicates cells without detected barcodes. The HERACLES tree was selected by fitted evolutionary likelihood, independently of the lentiviral barcodes. Numbers below each tree indicate the day-5 and day-7 integration counts after collapsing branches unsupported by recorder edits. **(d)** Minimum integration events required to explain the independent barcode distributions. Lower counts indicate greater agreement with the barcode annotations. HERACLES values show mean ± SD across three independent runs, with individual runs shown as points; baseline values represent single reconstructions. Dashed lines indicate the theoretical minimum of one integration per barcode.

HERACLES reconstructed lineages with integration-event counts comparable to LAML-Pro and substantially lower than the original PEtracer reconstruction (Fig. 6d). Across three independent runs, HERACLES required 49.3± 3.8 integrations on day 5 and 216.0 ±8.0 on day 7 (mean ± SD), compared with 49 and 220 for the available LAML-Pro tree, and 58 and 255 for the PEtracer tree. The HERACLES reconstruction shown in Fig. 6c was selected by its fitted evolutionary likelihood, without reference to the evaluation barcodes, and required 52 and 208 integrations, respectively. Thus, despite being 160x faster, HERACLES achieved similar agreement with the independent lineage markers to LAML-Pro across both integration rounds. Both substantially improved upon the original PEtracer reconstruction.

We next asked whether explicitly modeling a recorder’s editing mechanism could improve reconstruction accuracy. DNA Typewriter edits positions sequentially within a cassette, such that a downstream edit can occur only after preceding positions have been edited [2]. Treating sites independently ignores this ordering constraint. We therefore replaced the independent-site likelihood with a sequential cassette model while retaining the same hyperbolic optimization framework. Across 60 simulated datasets containing 250–1,000 cells and 12 five-site cassettes with missing data, the sequential model reduced mean rooted Robinson–Foulds (RF) distance by 0.081 relative to HERACLES with the independent-site model, improving 49 of 60 reconstructions (Supp. Fig. S12). It also reduced mean RF distance by 0.057 relative to neighbour joining on cassette-aware edit distances, improving all 60 reconstructions, and by 0.20–0.32 relative to the distance-based methods used in the original Typewriter study.

Together, these results show that HERACLES can accommodate distinct editing and missingness mechanisms within a common inference framework. Its PEtracer reconstructions agree with independent experimental lineage markers, while explicitly modeling the sequential constraints of DNA Typewriter improves reconstruction accuracy.

## 4 Discussion

HERACLES reconstructs cell lineages by optimizing hyperbolic coordinates under a composite likelihood of sampled quartets. By recovering rooted branching relationships directly from embedded distances, HERACLES avoids reconstructing a complete phylogeny during optimization. HERACLES can reconstruct lineages containing at least 150,000 cells, and we demonstrate improved accuracy over all existing methods at all tree sizes. HERACLES is also faster for massive datasets than all other methods except UPGMA, which it runs as a subroutine. Unlike sequential tree building methods, like LAML, HERACLES benefits from GPU optimization. Applications to KP-tracer and PEtracer produced trees consistent with independent lineage annotations, while adapting the likelihood to sequential editing improved reconstruction on simulated DNA Typewriter lineages.

The scalability of HERACLES relies on evaluating a small set of quartets rather than a complete phylogeny. Because quartet topologies and branch lengths are recovered directly from embedded distances, the choice of geometry is important. Euclidean embeddings of deeply branching trees incur increasing distortion, whereas hyperbolic space better preserves their hierarchical structure. Euclidean embeddings, as in VINE [17], avoid this distortion only by reconstructing a complete tree at each iteration, which imposes a valid phylogenetic structure before likelihood evaluation at a cost that grows at least quadratically with the number of cells. In contrast, HERACLES need only build a complete tree once at the end, although optionally building a tree at initialization can speed convergence.

The choice of quartets is also important. Neighbourhood-based sampling improved accuracy over uniform sampling, but does not guarantee coverage of every ancestral branch. Drawing neighbours within a fixed neighbourhood size may undersample branches near the root for larger trees; these branches that are separated by more edits and are easier to resolve at initialization. If necessary, future strategies could target poorly resolved ancestral relationships or improve coverage of inferred branches. Our quartet composite likelihood is an approximation to the likelihood of the full phylogeny. The latter considers each branch exactly once, and we hypothesize that ensuring that every branch is covered approximately uniformly improves the quality of the quartet composite likelihood approximation.

The computational cost of scoring quartets also depends on the evolutionary model. The star homoplasy assumptions used for conventional CRISPR recording restrict the possible ancestral states, making likelihood evaluation efficient. More general models could be incorporated using Felsenstein pruning, although considering additional ancestral states would increase the cost of scoring each quartet. Our results on PEtracer and DNA Typewriter demonstrate the benefit of adapting the likelihood to the recording mechanism. This flexibility could also allow other biological measurements, including transcriptional or spatial cell states, to inform lineage reconstruction through appropriate models of their relationship to ancestry.

Finally, HERACLES currently returns a single phylogeny without quantifying uncertainty. Extending the framework to optimize a variational distribution over cell coordinates [17–19], or to sample cell coordinates from a Gaussian approximation fitted to late-stage stochastic gradient descent iterates [32], could provide estimates of uncertainty in clade relationships and branch lengths through the resulting distribution of trees.

## Supplementary Information

### A Supplementary Methods

We provide the mathematical formulation and implementation details of HERACLES, beginning with the representation of lineage-recording data. The input is a character matrix *Y* ∈ ℤ^*n*×*S*^ of the observed states of *S* recording sites in *n* cells, in which *Y*_*is*_ = 0 denotes an unedited site, *Y*_*is*_ *>* 0 an edited allele and *Y*_*is*_ = −1 a missing observation, augmented with a synthetic, all-unedited root representing the ancestral state. The following sections describe the algorithm, recorder-specific likelihood models, simulated datasets and the evaluation procedures and metrics. Unless otherwise stated, all settings are the defaults of HERACLES v0.1.0.

#### A1 Initializing a hyperbolic representation of cell lineages

We estimate pairwise distances using the weighted Hamming distance of Cassiopeia [9]. Let *O*_*ij*_ = {*s* : *Y*_*is*_ ≠ −1, *Y*_*js*_ ≠ −1} denote the sites observed in both cells, including the synthetic founder as an additional cell. Then

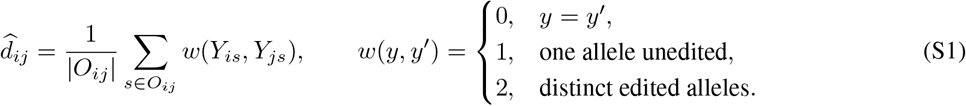

We set 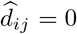 when *O*_*ij*_ is empty. The weights reflect the number of independent edits required under irreversible mutation. Sequential recorders instead use cassette-aware distances (Section A9). The resulting (*n* + 1) × (*n* + 1) matrix is used for embedding initialization and quartet sampling.

To initialize the embedding, we construct a preliminary tree from the estimated distances using neighbour joining [14], as implemented in RapidNJ [33], with negative branch lengths clamped to zero. The resulting tree-path distances provide a tree-consistent starting point even when the estimated distances are not additive. Initialization from UPGMA or Cassiopeia Greedy produces higher reconstruction error (Supp. Fig. S4). We obtain Euclidean coordinates from these distances using classical multidimensional scaling (MDS), which we favor over specialized hyperbolic embedding methods because the latter increased computational cost without improving reconstruction accuracy. Omitting the preliminary tree and applying MDS directly to the estimated distances also had little effect on final accuracy (Supp. Fig. S4).

Let Δ ∈ ℝ^*N* ×*N*^ denote the matrix of tree-path distances among the *N* = *n* + 1 cells, including the synthetic founder. Classical MDS constructs the centered Gram matrix

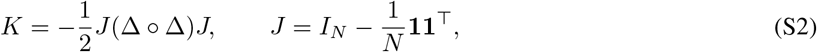

where *I*_*N*_ is the *N* ×*N* identity matrix, **1** ∈ ℝ^*N*^ is the vector of ones, and ° denotes elementwise multiplication. We obtain *D*-dimensional Euclidean coordinates by scaling the eigenvectors corresponding to the largest positive eigenvalues of *K* by the square roots of those eigenvalues. If fewer than *D* positive eigenvalues are available, the remaining coordinates are set to zero.

We rescale the Euclidean coordinates so that their largest absolute entry is 0.3, then map them onto the Lorentz hyperboloid using the exponential map at the origin. This initialization avoids large coordinate magnitudes that can cause numerical instability when evaluating hyperbolic distances through the Lorentz inner product, while allowing the embedding scale to be determined during likelihood optimization.

Finally, we apply a Lorentz boost, a distance-preserving transformation of the hyperboloid, to map the synthetic founder to the hyperbolic origin *ρ* = (1, 0, …, 0). Because Lorentz boosts preserve the Lorentz inner product, applying the same transformation to all cells leaves their pairwise hyperbolic distances unchanged. The founder is then held fixed throughout optimization, providing a common reference point for estimating ancestral depths.

The embedding dimension *D* controls the capacity of the representation to capture branching relationships, at a computational cost that grows linearly with *D* per optimization step. We use 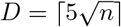, chosen from a sensitivity analysis showing diminishing improvements in reconstruction accuracy with increasing dimension (Supp. Fig. S8). The dimension remains fixed throughout optimization.

#### A2 Selecting informative quartets from edit-derived neighbourhoods

To preferentially sample quartets informative about local branching relationships, HERACLES draws all four cells of a quartet from a single neighbourhood of related cells. We estimate shared-ancestor depths using 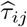, the clipped, root-referenced Gromov product of the edit-derived distances introduced in Section A1:

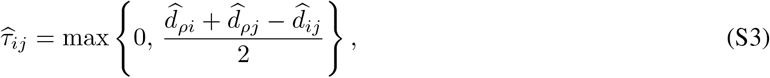

where *ρ* denotes the synthetic founder. For each cell *i*, we rank the other cells *j* by decreasing 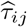, breaking ties by cell index, and keep the first *K* = min{*K*_max_, *n* − 1} ranks, where *n* is the number of observed cells, excluding the synthetic founder. These rankings are computed once and held fixed throughout optimization; updating them from the evolving embedding increased computational cost without improving reconstruction accuracy.

At each optimization step, HERACLES samples a batch 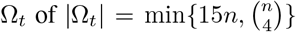 quartets. Each proposal consists of four distinct cells *a, b, c, d*, selected using the following procedure.

1. *Anchor*. Draw an anchor cell *a* uniformly from the *n* observed cells.
2. *Neighbourhood size*. Draw *u* uniformly from [0, 1) and set *k* = max {3, ⌊ (*K* + 1)^*u*^ ⌋}, so that *k* is approximately log-uniform on {3, …, *K*}. Log-uniform sampling gives equal probability to each doubling of neighbourhood size, favouring close relatives while retaining a chance of sampling more distant cells. *K* was set to 1024 in all experiments.
3. *Neighbours*. Draw three distinct ranks uniformly from {0, …, *k* −1} and take the corresponding cells in *a*’s ranking. The highest-ranked of the three becomes *b*, and the other two become *c* and *d*. Because all four cells lie within one neighbourhood of *a*, the quartet resolves a branching relationship at the depth set by *k*, without a separate outgroup window or a schedule over neighbourhood sizes.
4. *Acceptance*. Retain the proposal if

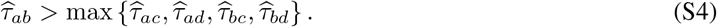

Because *b* outranks *c* and *d* in *a*’s ranking, 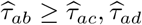 already holds; the test therefore requires that *b* is also_*ab ac ad*_ closer to *a* than to *c* or *d*, and rejects ties. This favours quartets with a closely related pair without constraining the topology subsequently inferred from the hyperbolic embedding.

Rejected proposals are replaced by additional draws. Each round generates three proposals per unfilled batch position; after eight rounds, any remaining positions are filled by uniform sampling. If all estimated shared-ancestor depths are zero, no proposal passes the acceptance test and sampling is therefore uniform.

#### A3 Recovering rooted quartets directly from hyperbolic distances

HERACLES recovers the rooted topology and branch lengths of each sampled quartet directly from hyperbolic distances, without explicitly representing ancestral nodes in the embedding. In a rooted tree, the depth of the most recent common ancestor (MRCA) of two cells is determined by their distances to the root and to one another. We approximate this depth using the root-referenced Gromov product

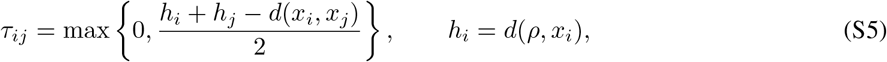

where *x*_*i*_ denotes the hyperbolic coordinate of cell *i, ρ* is the fixed founder, and *h*_*i*_ is the root-to-leaf distance. For additive tree distances, the Gromov product recovers ancestral depth exactly; for hyperbolic distances, it provides an approximation. Unlike the edit-derived estimates 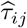 used for quartet sampling, these depths change with the embedding during optimization.

**Figure S1:**
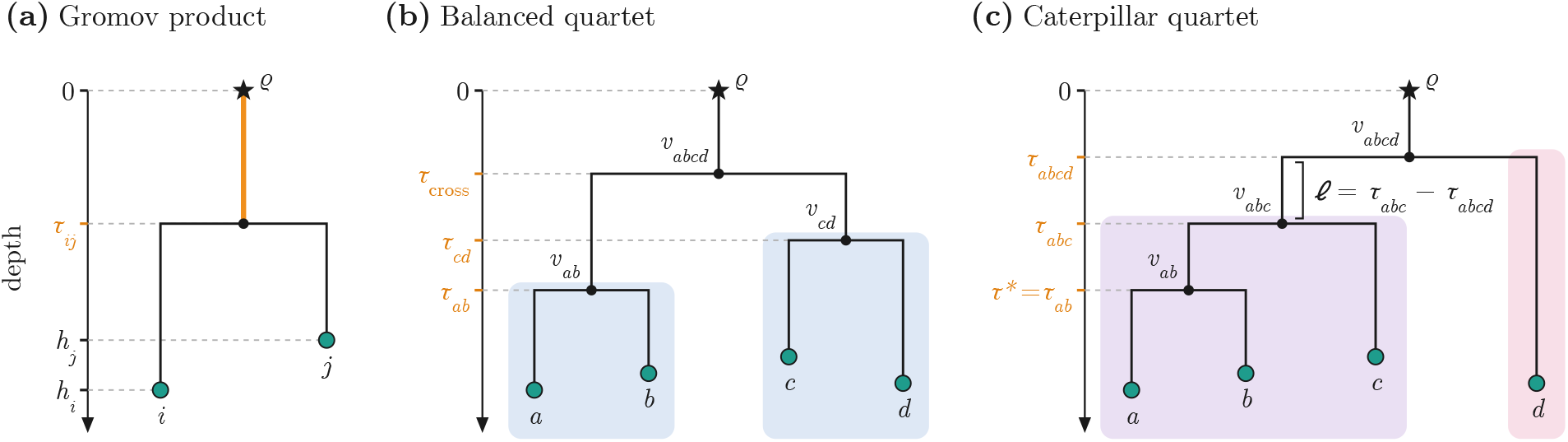
Recovering ancestral depths and branch lengths from hyperbolic distances. Depth increases downward from the founder *ρ*. **(a)** The root-referenced Gromov product *τ*_*ij*_ estimates the depth of the MRCA of two sampled cells. **(b)** In a balanced quartet, *τ*_cross_ locates the first split, while *τ*_*ab*_ and *τ*_*cd*_ locate the two cherries. **(c)** In a caterpillar quartet, *τ*_*abcd*_, *τ*_*abc*_ and *τ*_*ab*_ locate successive ancestral nodes. Branch lengths are differences between inferred node depths, illustrated by *l* = *τ*_*abc*_ − *τ*_*abcd*_.

#### Topology and rooting

For each sampled quartet *Q* = {*a, b, c, d*}, we first identify its unrooted topology using the four-point condition, computing

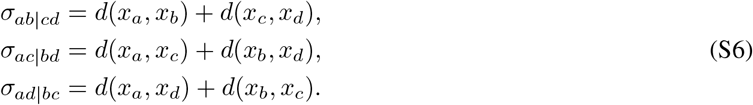

We select the split with the smallest sum, resolving ties in the order listed.

To determine the rooted shape, consider the selected split *ab*|*cd* and define the mean cross-pair depth

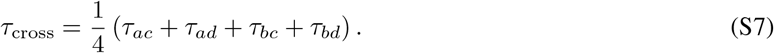

If *τ*_*ab*_≥ *τ*_cross_ and *τ*_*cd*_≥ *τ*_cross_, the quartet is *balanced*, with topology ((*a, b*), (*c, d*)). The first common ancestor of all four cells is at depth *τ*_cross_.

Otherwise, the quartet is a *caterpillar*. The outgroup is selected from the pair whose within-pair Gromov product falls below *τ*_cross_, choosing the cell with the smaller mean Gromov product to the opposite pair. Among the remaining three cells, the cherry is the pair with the largest Gromov product. We relabel the cells after topology selection so that the caterpillar is written as (*d*, (*c*, (*a, b*))), where *d* is the outgroup, *c* branches next, and {*a, b*} forms the cherry. The nested triple’s cherry is determined from its own Gromov products and need not coincide with the pair implied by the initial four-point split.

##### Branch lengths

Although only sampled cells have embedding coordinates, their Gromov products provide estimates of the depths of unobserved ancestral nodes. We denote an internal node by *v*_*A*_, where *A* is its descendant cell set, and its inferred depth by *τ*_*A*_. Thus, *v*_*abcd*_ is the MRCA of all four sampled cells, while *v*_*ab*_ is the MRCA of the cherry {*a, b*} (Supp. Fig. S1).

For a balanced quartet ((*a, b*), (*c, d*)), the internal node depths are

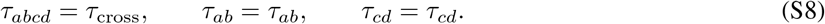

Here *τ*_cross_ estimates the depth of *v*_*abcd*_, while the within-pair Gromov products give the depths of *v*_*ab*_ and *v*_*cd*_.

For a caterpillar (*d*, (*c*, (*a, b*))), the three successive ancestral depths are

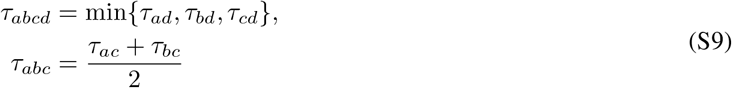

These correspond to the depths of *v*_*abcd*_, where the outgroup *d* separates; *v*_*abc*_, where *c* separates from the cherry; and *v*_*ab*_, the MRCA of *a* and *b*, respectively.

For either topology, the depth of the founder is zero and the depth of each sampled cell *i* is *h*_*i*_. Writing depth(*q*) for the depth of any node *q*, the length of an edge from parent *p* to child *q* is

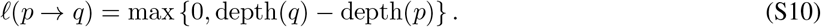

Negative differences are clamped to zero to ensure nonnegative branch lengths when embedded distances are not exactly tree-like.

#### A4 Modeling CRISPR editing and missing observations

The rooted quartets recovered in Section A3 are evaluated under a probabilistic model of recording-state evolution. Here we describe the continuous-time Markov chain (CTMC) used for conventional CRISPR/Cas9 recording; extensions for other recording technologies are given in Section A9.

##### CRISPR/Cas9 mutation model

We follow the probabilistic mixed-type missing (PMM) model of LAML [11], which accounts for irreversible editing, heritable deletion and stochastic observation dropout. Each recording site *s* evolves independently, beginning in the unedited state 0 and acquiring allele *α* ∈ {1, …, *K*_*s*_} at rate *λq*_*sα*_, where *λ* is the editing rate, *K*_*s*_ is the number of possible edited alleles at site *s*, and *q*_*sα*_ is the probability of acquiring allele *α* upon editing, with 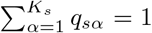. Once edited, a site cannot revert or acquire a different allele, although identical alleles may arise independently in different lineages.

Missing observations arise through two mechanisms. Heritable deletion occurs at rate *ε*_*H*_ from any nondeleted state and is inherited by descendants, producing correlated missing observations within a clade. Stochastic dropout independently obscures surviving states in sampled cells with probability *ε*_*S*_.

Editing and deletion together define a CTMC on the latent states {0, 1, …, *K*_*s*_, ∅}, where ∅ denotes the deleted state (Fig. 2c). Its rate matrix is

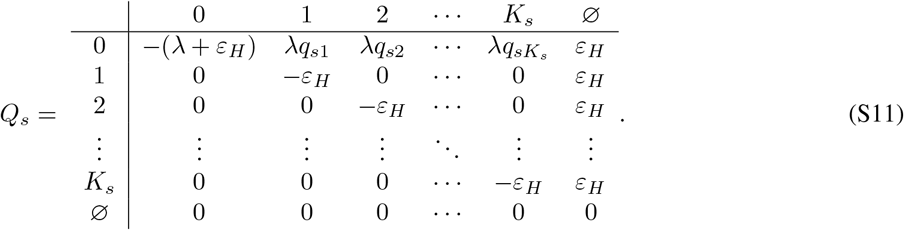

The transition probabilities along a branch of length *l* are given by 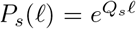, with *P* (*z*→ *z*^′^ | *l*) denoting the probability of transitioning from state *z* to state *z*^′^. Because deletion occurs at the same rate from every nondeleted state, survival along a branch is independent of the editing process, with probability 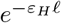. For conventional Cas9 recording, we fix *λ* = 1, expressing branch lengths in units of the editing timescale.

Let *z*_*vs*_∈ {0, 1, …, *K*_*s*_, ∅} denote the latent state of site *s* at node *v*, including unobserved ancestral nodes. For a sampled cell *i*, the observation model is

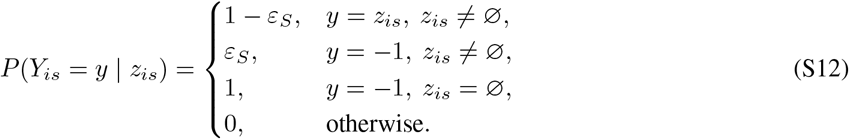

Thus, heritable deletion and stochastic dropout both produce the observed missing state −1, but only deletion is inherited along the lineage.

##### Parameters and priors

The allele probabilities *q*_*sα*_, deletion rate *ε*_*H*_ and dropout probability *ε*_*S*_ are estimated jointly with the embedding. Allele probabilities are parameterized by softmax-transformed logits, initialized uniformly and learned without a prior, either as a separate distribution for each site or as a single distribution shared across sites. The deletion rate and dropout probability are optimized on unconstrained scales, 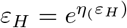 and 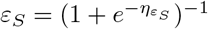.

Both missingness parameters have priors with modes at their initial values: a Gamma prior on *ε*_*H*_ with mode 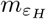 and variance 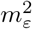, and a Beta prior on *ε*_*S*_ with mode 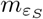 and variance 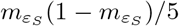. Their combined negative log-prior, normalized by (|Ω_*t*_| + *n*)*S*, defines ℒ _prior_.

Allele probabilities are initialized uniformly over the specified allele support at each site, and both missingness parameters are initialized at 0.1 by default. Dataset-specific initialization settings are described in the corresponding experimental methods.

#### A5 Computing quartet likelihoods

For each sampled quartet *Q*, HERACLES evaluates the probability of its observed recording states under the rooted topology and branch lengths inferred in Section A3. We use Felsenstein’s pruning recursion [30] to marginalize over the unobserved ancestral states, with the founder fixed in the unedited state. The likelihood is assembled from smaller subtrees, exploiting the irreversibility of CRISPR editing to reduce the number of ancestral states that must be considered (Supp. Fig. S2).

**Figure S2:**
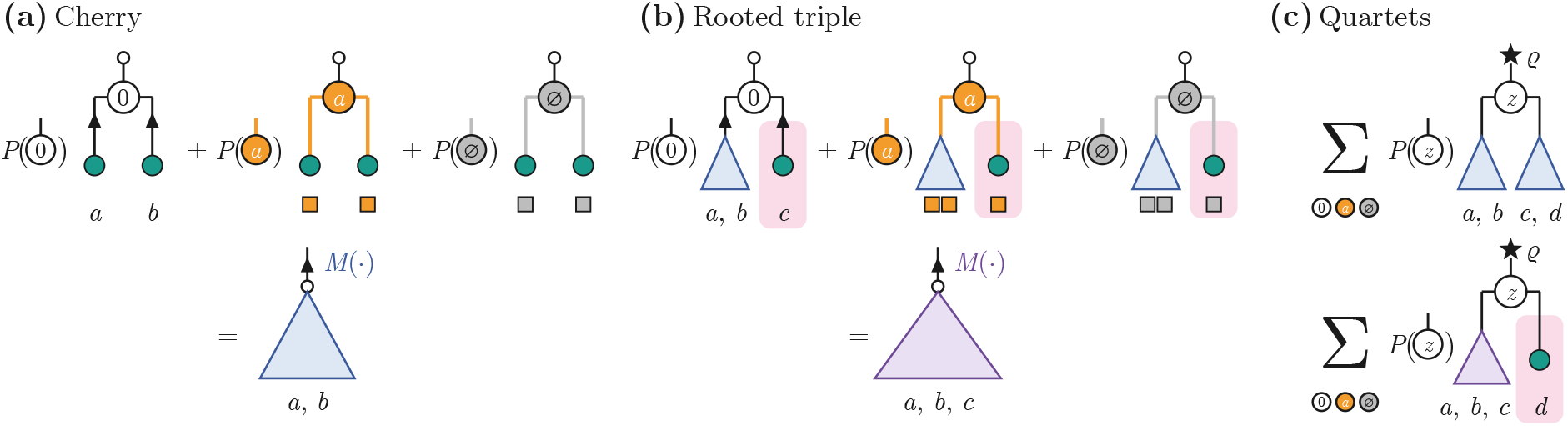
Recursive computation of quartet likelihoods. **(a)** A cherry likelihood sums over possible ancestral states. **(b)** A rooted triple combines a cherry with a third cell. **(c)** Balanced and caterpillar quartets combine two cherries or a triple and an outgroup, respectively. Ancestral states are weighted by transition probabilities along the connecting branches; triangles represent previously evaluated subtrees.

##### Pruning recursion

Felsenstein pruning computes the likelihood from the leaves upward, combining the contributions of progressively larger subtrees. For a recording site *s*, let *F*_*v*_(*z*_*v*_) denote the probability of the observations in the subtree rooted at node *v*, conditional on *v* being in state *z*_*v*_. At a sampled cell *i*, this is simply the observation probability defined in Eq. (S12):

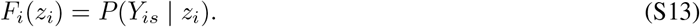

To incorporate a subtree into the likelihood at its parent, we must account for evolution along the connecting branch. Let *p* be the parent of *v*, and let *l*_*pv*_ denote the length of the branch *p* →*v*. Given the parent’s state *z*_*p*_, the state at *v* is unknown, so we sum over its possible values, weighting the likelihood of the subtree by the corresponding transition probabilities:

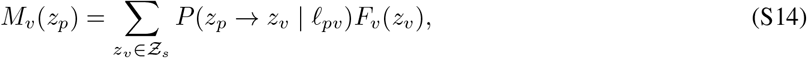

where Ƶ_*s*_ = {0, 1, …, *K*_*s*_, ∅} is the set of latent recording states. The resulting message *M*_*v*_(*z*_*p*_) gives the probability of the observations in the subtree rooted at *v*, conditional on the state of its parent, accounting for evolution along the connecting branch.

At an internal node *v*, each child *c* contributes a message *M*_*c*_(*z*_*v*_) that incorporates the observations in its subtree and the transition probabilities along the branch *v* →*c*. Given the state *z*_*v*_, observations in different child subtrees are independent, so their probabilities multiply:

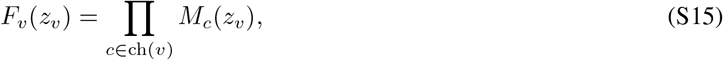

where ch(*v*) denotes the children of *v*. This produces the likelihood of all observations below *v*, conditional on its state. The same two operations are repeated toward the founder to obtain the likelihood of the complete quartet.

For a balanced quartet ((*a, b*), (*c, d*)), the two cherry likelihoods at *v*_*ab*_ and *v*_*cd*_ are combined at *v*_*abcd*_. For a caterpillar (*d*, (*c*, (*a, b*))), the cherry at *v*_*ab*_ is first combined with cell *c* at *v*_*abc*_, and the resulting triple is combined with outgroup *d* at *v*_*abcd*_. The recursion terminates at the quartet root *v*_*abcd*_, whose state is unobserved and is distinct from the founder *ρ*. The stem connecting them represents the shared evolutionary history before the four sampled cells diverge. Edits or heritable deletions acquired along this stem may be inherited by all four descendants and must therefore be included in the likelihood. Since the founder is fixed in the unedited state, we marginalize over the possible states at *v*_*abcd*_:

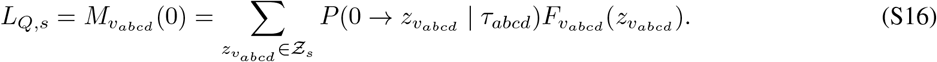

##### Efficient evaluation under irreversible editing

The transition model in Section A4 restricts the ancestral states that can contribute to the pruning recursion. Once a node carries an edited allele *α*, all surviving descendants must retain that allele; if the node is deleted, all descendants are missing. Consequently, an edited ancestral state has nonzero partial likelihood only if all observed descendant alleles agree.

Let *e*(*l*) = *e*^−*λl*^ and 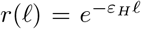 denote the probabilities of no editing and no deletion, respectively, along a branch of length *l*. Applying the CTMC transitions to Eq. (S14) gives

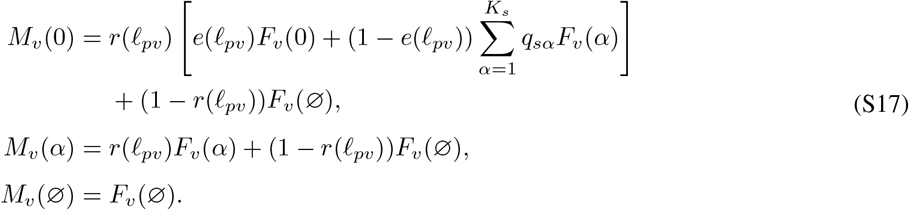

Here *F*_*v*_(∅) is nonzero only when all observations descended from *v* are missing, since deletion is irreversible.

The sum over edited ancestral states can be evaluated without enumerating all *K*_*s*_ alleles. If any descendant is observed as unedited, or if two distinct edited alleles are observed, every edited ancestral state has zero likelihood. If the observed descendants carry a single edited allele *α*^*^, only that allele can contribute. When all descendants are missing, the partial likelihood is identical for every edited ancestral state, and the weighted sum simplifies using ∑ _*α*_ *q*_*sα*_ = 1.

Thus, only a constant number of likelihood quantities need to be evaluated per node, and computing the likelihood of a quartet requires *O*(*S*) operations across *S* recording sites.

##### Root-to-leaf likelihood

In addition to the quartet likelihoods, HERACLES evaluates the probability of each cell’s observed state along its direct path from the unedited founder. For a cell at depth 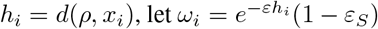 denote the probability that its recording site survives deletion and is observed. The resulting probabilities are

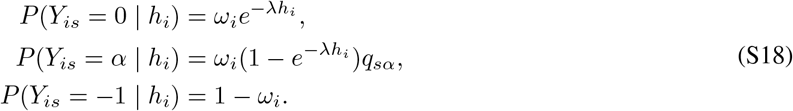

These root-to-leaf likelihoods provide information about absolute evolutionary depth that may be weakly constrained by sampled quartet relationships alone.

#### A6 Optimizing the embedding with a regularized composite likelihood

HERACLES optimizes the hyperbolic embedding using a regularized composite likelihood of sampled quartets and root-to-leaf paths. The founder terms constrain evolutionary depths, while geometric penalties encourage tree-like distances and, optionally, approximately equal root-to-leaf depths.

##### Composite objective

At optimization step *t*, let Ω_*t*_ denote the sampled quartets, *L*_*Q,s*_ the likelihood of quartet *Q* at site *s*, and *L*_*ρ,i,s*_ the root-to-leaf likelihood defined in Section A5. The total loss is

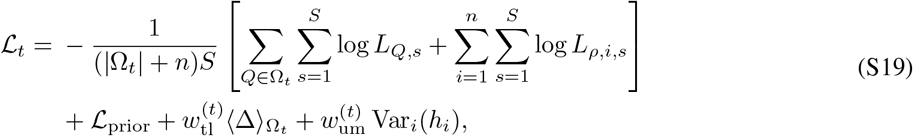

where ℒ_prior_ is the normalized negative log-prior defined in Section A4, 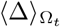 measures departures from tree-additive distances, and Var_*i*_(*h*_*i*_) measures variation in root-to-leaf depths. Each quartet-site and founder-path-site likelihood receives equal weight, without an additional mixing parameter.

##### Tree-like penalty

Hyperbolic distances need not satisfy the four-point condition of an additive tree metric. For each sampled quartet *Q*, let *σ*_(1)_(*Q*) ≥*σ*_(2)_(*Q*) ≥*σ*_(3)_(*Q*) denote its three ordered four-point distance sums (Section A3). We define

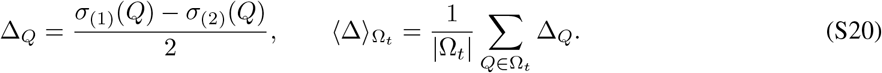

An additive tree metric has Δ_*Q*_ = 0 for every quartet. Unlike classical Gromov *δ*-hyperbolicity, which is determined by the maximum four-point defect, we penalize the mean defect over the sampled batch, reusing the quartets evaluated for the likelihood. Because strong tree-like regularization restricts exploration of alternative branching relationships, we optimize in two stages: initially without this penalty, then with its weight adjusted adaptively.

##### Ultrametric penalty

For contemporaneously sampled cells evolving at approximately constant rates, root-to-leaf depths are expected to be similar. We therefore optionally penalize their variance,

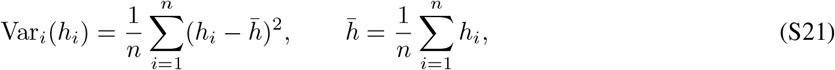

where *h*_*i*_ = *d*(*ρ, x*_*i*_) is the depth of cell *i*. This encourages approximately ultrametric distances without enforcing identical depths. The penalty may be disabled when cells are sampled asynchronously or editing rates vary across lineages; it was enabled in all analyses reported here.

##### Adaptive penalty weights

Rather than tuning fixed coefficients, HERACLES adjusts the penalty weights to target a mean four-point defect *τ*_tl_ = 0.01 and a root-to-leaf depth variance equal to 75% of its initial value:

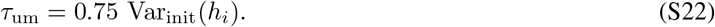

After each optimization step, the active weights are updated using a dual-ascent-inspired rule:

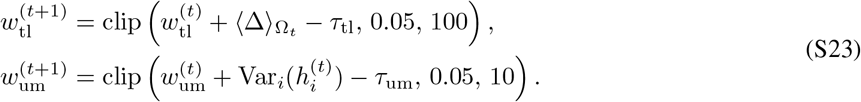

Here clip(*x, a, b*) restricts *x* to [*a, b*]. Both weights start at zero and are subject to the stated bounds once their controllers are active. The effect of each penalty is shown in Supplementary Fig. S6.

##### Coordinate optimization

We jointly optimize the embedding coordinates and the learnable parameters of the evolutionary model using Riemannian Adam [31], as implemented in geoopt v0.5.1 [34]. Although quartet topology selection is discrete, branch lengths are differentiable almost everywhere with respect to the embedding coordinates within regions of fixed topology. Gradients therefore propagate through the inferred branch lengths, allowing continuous optimization to change the reconstructed quartet relationships.

We use a constant learning rate of 0.02 and evaluate the unpenalized composite loss on a fixed validation batch every 100 steps. Each stage terminates after three consecutive relative changes below 10^−4^, with defaults of a minimum of 1,500 and a maximum of 2,500 steps.

#### A7 Reconstructing the final phylogeny

Following optimization, HERACLES reconstructs a phylogeny from the pairwise hyperbolic distances between sampled cells and the synthetic founder. By default, we use RapidNJ [33], which accelerates conventional neighbour joining by sorting distances and pruning unnecessary pair comparisons while preserving the exact neighbour-joining criterion. The resulting tree is rooted on the edge connecting the founder, which is then removed; neighbour-joining branch lengths are retained.

Neighbour joining exactly reconstructs additive tree metrics and is guaranteed to recover the correct topology when distance errors are less than half the shortest edge length [16]. HERACLES encourages tree-additive distances through geometric regularization, while the evolutionary likelihood constrains their agreement with the observed recording data. Although these objectives do not guarantee exact recovery, they provide a principled basis for distance-based reconstruction. For very large datasets, we instead use UPGMA [15], which admits efficient quadratic-time reconstruction and is appropriate for approximately ultrametric distances (Section A8).

#### A8 Reconstructing atlas-scale lineages

For simulations of 25,000–150,000 cells (main Fig. 4c and Supp. Fig. S11), we adapted the HERACLES implementation to reduce computation time and memory requirements. At these scales, constructing dense pairwise distance matrices, ranking all potential neighbours and reconstructing an initial tree become increasingly expensive. We therefore use landmark-based initialization, GPU-accelerated neighbourhood construction, smaller quartet batches and a faster final tree read-out. The evolutionary likelihood, neighbourhood-based sampling strategy and optimization procedure remain as described above. During optimization, no full *n*× *n* distance matrix is stored; the final tree reconstruction remains the principal memory bottleneck. Coordinates and likelihood calculations use double precision throughout.

##### Initialization

We use landmark multidimensional scaling (MDS) [35], omitting the preliminary neighbour-joining tree to reduce initialization time. We select *m* = max {500, 2*D* + 1} landmark cells with a fixed seed, including the founder, and compute their weighted-Hamming distances to all cells. Classical MDS embeds the landmarks in *D* dimensions, and a Nyström extension places the remaining cells using their squared distances to the landmarks. This requires an *m*× *n* rather than an *n* ×*n* distance matrix.

The centred landmark Gram matrix has rank at most *m* −1, so *m > D* is necessary to recover *D* dimensions. Because weighted-Hamming distances need not be Euclidean, some eigenvalues may also be negative or near zero. We therefore use approximately 2*D* landmarks, with a minimum of 500, to provide a margin of usable dimensions. Eigenvalues below 10^−8^ of the largest are discarded, and unused dimensions are initialized at zero (Section A1). The resulting coordinates are mapped to the hyperboloid as in the standard initialization.

##### Sampling neighbourhoods

Neighbourhood-based sampling ranks potential relatives by estimated common-ancestor depth (Supp. Methods A2). Rather than storing all *n*(*n*− 1) ranked indices, we retain the 1,024 nearest neighbours of each cell, which is the largest neighbourhood the sampler can draw, so the truncation does not alter quartet sampling. We construct these neighbourhoods using GPU matrix multiplication to improve runtime. Let *V, E* and *Z* be *n* × *S* indicators of observed, edited and unedited sites, and let *H* be the one-hot encoding of edited alleles. The matrix *V V* ^⊤^ counts jointly observed sites, while

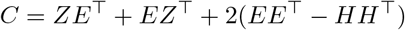

counts unedited–edited mismatches once and distinct edited alleles twice. Thus 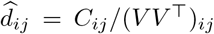 for jointly observed pairs, and zero otherwise (Eq. (S1)).

Matrix products are evaluated on the GPU in blocks of 1,024 rows using single precision, which exactly represents the integer counts. Each block is converted to estimated common-ancestor depths 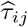 and reduced to its top 1024 neighbours per cell. This avoids materializing the full matrix and reduces ranking storage to *O*(*n*). Common-ancestor depths needed for quartet acceptance outside the stored neighbourhoods are recomputed directly from the recording data on the GPU.

##### Quartet optimization

We use 8*n* quartets per step rather than the default 15*n*, reducing likelihood-evaluation time. Batches are divided into micro-batches of at most 250,000 quartets, with gradients accumulated before each parameter update to preserve the sampled objective while limiting GPU memory use. Optimization otherwise uses the standard Riemannian Adam settings and convergence criteria.

##### Tree reconstruction

We decode the optimized embedding using average-linkage UPGMA, which is *O*(*n*^2^) and faster than neighbour joining. Founder-referenced Gromov products are converted to dissimilarities

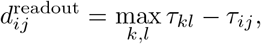

so that deeper shared ancestry corresponds to smaller dissimilarities. Dissimilarities are computed in blocks and written to a condensed double-precision vector of *n*(*n*−1)*/*2 entries. This final read-out determines peak host-memory use; at 150,000 cells, the vector alone occupies approximately 90 GB.

###### Computational resources

GPU reconstructions used eight CPU cores and one NVIDIA H100, with 48–240 GB of host memory requested; CPU-only comparisons used 16 cores. Peak resident memory ranged from 7 GB at 25,000 cells to 190 GB at 150,000 cells, including tree reconstruction. Runtime and memory results are reported in Supp. Fig. S11.

#### A9 Recorder-specific likelihoods

As lineage-recording technologies become increasingly diverse, their distinct editing dynamics and missing-data mechanisms present a challenge for methods tailored to individual recorders. HERACLES separates the evolutionary likelihood from the underlying inference framework, allowing recorder-specific models to be incorporated without redesigning the algorithm. Here, we demonstrate this flexibility with two extensions that modify the quartet and root-to-leaf likelihoods while preserving the embedding optimization, quartet sampling, regularization, and tree reconstruction.

##### PEtracer

PEtracer [3] uses genomically integrated lineage-tracing cassettes, each carrying an integration barcode and three independently edited sites. Because the three sites of a cassette are read out together, a failure to recover the cassette, or stable silencing of the integrated cassette, removes all three sites at once. Consistent with this architecture, missingness in the PEtracer dataset was entirely cassette-wide: of 37,296 cell–cassette observations (3,108 cells across 12 cassettes), 4,562 (12.2%) were entirely missing, and none had only one or two missing sites.

We therefore model stochastic dropout and heritable silencing at the cassette rather than the site level, while retaining independent editing dynamics at individual sites. Because both mechanisms remove the sites of a cassette together, the conventional likelihood counts the same missingness pattern once per site. We correct this as

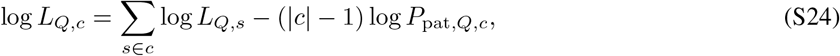

where *P*_pat,*Q,c*_ is the probability of the cassette’s missingness pattern, evaluated with editing disabled. This counts the shared missingness contribution once while retaining the edit-state likelihoods of individual sites. The same correction is applied to root-to-leaf likelihoods. A cassette-specific dropout rate *ε*_*S,c*_ can accommodate differences in detection efficiency among integrated cassettes. As the observed missing fraction per cassette ranges from 1.0% to 31.5% (about 30-fold) the PEtracer analysis used a single heritable silencing rate shared across cassettes and a separate dropout rate for each cassette.

##### Sequential editing

Sequential lineage recorders such as DNA Typewriter [2] accumulate edits in a prescribed order within each cassette. Unlike conventional Cas9 recorders, in which sites mutate independently, an edit at position *j* can occur only after positions 1, …, *j* −1 have been edited (Fig. S3a). Each cassette therefore records an ordered sequence of mutations, requiring a joint evolutionary model of its recording sites. We replace the independent-site likelihood with a cassette-level continuous-time Markov chain (CTMC) in which each editing event extends an existing edited prefix by one position.

For a cassette of *L* positions, the state consists of the number of edited positions *c* ∈ {0, …, *L*} and their ordered alleles (*a*_1_, …, *a*_*c*_). The first edit occurs at rate *αλ* and subsequent edits at rate *λ*, with each new allele drawn independently from distribution *q*. Thus, *λ* controls the rate of cassette extension, while *α* allows initiation to occur at a different rate. The fully edited state *c* = *L* is absorbing.

**Figure S3:**
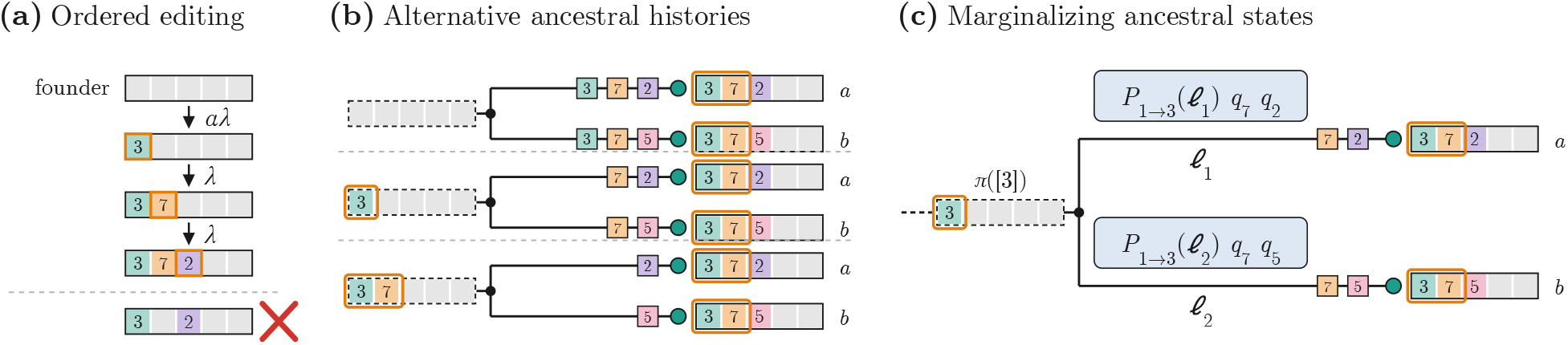
Modeling sequential lineage recording. **(a)** Edits extend an ordered prefix, with initiation rate *αλ* and continuation rate *λ*. **(b)** Alternative ancestral prefixes compatible with the same observed descendants. **(c)** Quartet likelihoods marginalize over compatible ancestral states, multiplying branch transition probabilities determined by edit counts, allele identities and branch lengths.

The sequential constraint restricts the ancestral states compatible with the observed cells. An ancestral cassette can contain only a prefix of its descendants’ edits, preserving their allele identities and order. For example, cells carrying (3, 7, 2) and (3, 7, 5) may descend from an ancestor with no edits, the prefix (3) or the prefix (3, 7) (Fig. S3b). These alternatives correspond to different histories of inherited and independently acquired edits. The likelihood therefore marginalizes over compatible ancestral prefixes rather than assigning a single ancestral state.

###### Transition probabilities

Let *P*_*ij*_(*l*) denote the probability that a cassette progresses from *i* to *j* edited positions along a branch of length *l*. When initiation and continuation occur at the same rate (*α* = 1), editing follows a Poisson process until the cassette is saturated. For *i* ≤ *j < L*,

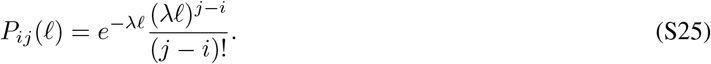

The probability of reaching the saturated state *L* is the remaining probability mass.

When *α* ≠ 1, only transitions starting from an unedited cassette differ. To reach state *j*, the first edit must occur at some time *t* along the branch, followed by exactly *j* − 1 continuation events during the remaining time *l* − *t*. Marginalizing over the initiation time gives, for 1 ≤ *j < L*,

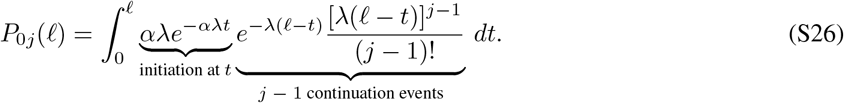

The probability of remaining unedited is *P*_00_(*l*) = *e*^−*αλl*^, and the probability of reaching saturation is 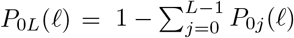. For computational efficiency, we evaluate the initiation-time integral using an analytic recurrence rather than numerical quadrature. Where the recurrence is numerically unstable, we instead obtain the transition probabilities by exponentiating the CTMC generator.

These probabilities describe the number of edits accumulated along a branch. The probability of a particular allele sequence additionally depends on the identities of the new edits. For a transition extending the prefix (*a*_1_, …, *a*_*i*_) to (*a*_1_, …, *a*_*j*_),

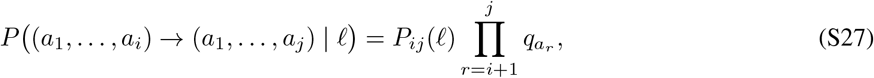

provided the ancestral prefix is preserved; all other transitions have probability zero.

###### Quartet likelihood

We calculate cassette likelihoods using the pruning recursion defined in Section A5, replacing the independent-site states and transition probabilities with ordered cassette prefixes. As in the conventional model, *F*_*v*_(*z*_*v*_) denotes the probability of observations in the subtree rooted at *v* conditional on its state *z*_*v*_, and *M*_*v*_(*z*_*u*_) includes the transition from its parent *u*:

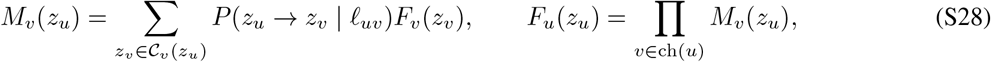

where *C*_*v*_(*z*_*u*_) contains the descendant prefixes compatible with the ancestral state and the observed edits. The founder is fixed in the unedited state, and the same transition model is used for root-to-leaf likelihoods. By restricting ancestral states to compatible prefixes, the recursion avoids enumerating arbitrary combinations of edited sites (Fig. S3c).

###### Missingness and parameter estimation

As in the conventional model (Section A4), we distinguish heritable deletion from stochastic observation dropout. A cassette can be permanently deleted at rate *ε*_*H*_ along any branch, after which it remains missing in all descendants. In contrast, stochastic dropout obscures an otherwise surviving cassette at a sampled cell with probability *ε*_*S*_, without affecting its descendants or relatives. Both mechanisms act on entire cassettes rather than individual recording positions, so a missing cassette is represented by a single missing observation in the likelihood. Partially observed cassettes are treated as entirely missing during inference.

Each cassette log-likelihood is divided by *L* to retain the per-site scale of the conventional HERACLES objective. The continuation rate *λ* is estimated jointly with the embedding, together with the allele probabilities, deletion rate *ε*_*H*_ and dropout probability *ε*_*S*_; the initiation ratio was fixed at *α* = 1 in all analyses reported here.

###### Prefix-based distances

Sequential editing also motivates a distance based on shared edited prefixes. For cells *i* and *j* and cassette *k*, let *c*_*ik*_ and *c*_*jk*_ denote their numbers of edited positions, and let *p*_*ijk*_ be the length of their longest identical edited prefix, ending at the first differing allele or unedited or missing position. We define the cassette-aware distance

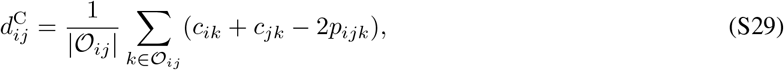

where *O*_*ij*_ is the set of cassettes observed in both cells. This counts edits not shared by the two prefixes, averaged over jointly observed cassettes. It ranges from 0 for identical prefixes to 2*L* for fully edited cassettes differing at the first position, and reduces to the conventional weighted-Hamming distance when *L* = 1.

For comparison, the original DNA Typewriter study [2] used a distance based only on shared edits,

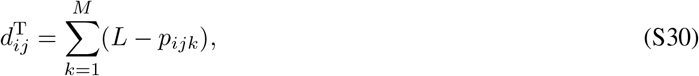

where *M* is the number of cassettes. Unlike *d*^C^, this distance ignores edits accumulated beyond the shared prefix and assigns a contribution of *L* even when both cassettes are unedited. It also sums over all cassettes rather than averaging over jointly observed ones. Because the original study retained only cells with fully recovered cassettes, it did not specify a missing-data rule; for our comparisons, a missing position terminates the shared prefix and contributes *L* for that cassette. HERACLES embeddings are initialized by MDS on *d*^C^.

#### A10 Simulation datasets and experimental design

We evaluated HERACLES on published LAML and Startle benchmarks, additional Cassiopeia-generated lineages spanning 1,000–150,000 cells, and sequential-editing simulations. Separate development collections were used for ablations and computational evaluation.

##### Published lineage-reconstruction benchmarks

The LAML benchmark [11] comprises five missingness regimes at 250 cells and 30 recording sites, with nominal heritable/stochastic missingness proportions of 0/100, 25/75, 50/50, 75/25 and 100/0. Three additional conditions contain 500, 750 and 1,000 cells under the 50/50 regime. Each of the eight conditions contains 50 datasets spanning ten allele-alphabet settings and five subsamples of a simulated 1,024-leaf tree. The Startle benchmark [10] comprises 1,736 datasets across 136 configurations with 50–200 cells, 10, 20 or 30 recording sites, per-edge editing probabilities of 0.1, 0.2 or 0.3, stochastic dropout rates of 0–0.2, 25 alleles and no heritable deletion. The parameter grid is incomplete, with 8–21 replicates per configuration. For both benchmarks, we used the published character matrices and ground-truth trees without regenerating the simulations.

##### Scalability simulations

We generated lineages using Cassiopeia’s birth–death–fitness tree simulator (birth waiting-time scale 0.5, death waiting-time scale 1.5, no fitness effects) and Cas9 recording simulator [9]. For 1,000, 2,000, 5,000 and 10,000 cells, we varied recording sites (30 or 60), mutation rate (0.1, 0.2 or 0.3) and heritable silencing rate (0.02 or 0.03), with stochastic dropout fixed at 0.2 and applied using an independent random stream. Each site had 30 possible alleles with exponentially sampled priors (scale 10^−5^). Five replicates per parameter combination yielded 60 datasets per size.

Atlas-scale simulations comprised five replicates each at 25,000, 50,000, 100,000 and 150,000 cells, with 60 sites, mutation rate 0.2, heritable silencing rate 0.02 and stochastic dropout 0.2. Cassiopeia was pinned to commit `1ee5959e` (v2.1.0).

##### Sequential-recording simulations

We generated DNA Typewriter-like recordings using Cassiopeia’s sequential lineage-recording simulator, with 12 cassettes of five ordered editing positions and 30 alleles with exponentially sampled priors (scale 10^−5^). We varied cell count (250, 500 or 1,000) and initiation/continuation rates ((0.5, 0.5), (1.0, 0.5), (0.5, 0.1) or (0.2, 1.0)), with heritable silencing 0.01 and stochastic dropout 0.05. Five replicates per condition yielded 60 datasets. We compared the sequential and independent-site HERACLES likelihoods with distance-based reconstruction using cassette-aware and shared-prefix distances.

We evaluated six reconstruction methods on this data. Two were HERACLES variants, one using the sequential cassette likelihood described above and the other using the conventional independent-site likelihood. Both used the same optimization settings as the other HERACLES experiments: Riemannian Adam with a constant step size of 0.02, convergence checked every 100 steps, and termination after three consecutive relative loss changes below 10^−4^. Both missing-data rates were initialized at 0.1, so that only the evolutionary likelihood differed between variants.

The remaining four methods were neighbour joining and UPGMA, each applied to the cassette-aware distance *d*^C^ (Eq. (S29)) and the Typewriter distance *d*^T^ (Eq. (S30)). The cassette-aware baselines used the same distances as HERACLES initialization, isolating the contribution of likelihood-based optimization, while the Typewriter-distance baselines reproduced the original study’s distance-based reconstruction methods with our missing-data extension. All inferred and true trees were contracted under Camin–Sokal parsimony using each replicate’s character matrix and evaluated by collapsed rooted Robinson–Foulds (RF) distance (Section A12).

##### Evaluating algorithmic components

Objective-order and quartet-sampling ablations used two collections of 500 cells and 30 sites (Table S1): a stochastic-missingness collection of 25 replicates and the mixed regime of the development collection (20 replicates). Pairwise, triplet and quartet objectives used uniform sampling under matched initialization, regularization, parameter learning and reconstruction settings. The all-pairs objective evaluated every cell pair, the sampled-pairs objective sampled 30*n* pairs per step, the triplet objective sampled 20*n* triples per step, and the quartet objective sampled 15*n* quartets. Neighbourhood-based and uniform quartet sampling were then compared under otherwise identical settings. All ablation fits, including those without the geometric penalties, used a maximum budget of 20,000 steps. Separate collections assessed initialization, embedding dimension and computational performance (Table S1); missing-data parameter recovery and the four-point defect were assessed on the 250-cell conditions of the LAML benchmark.

**Table S1:** Simulation collections for algorithmic evaluation. All collections use *µ* = 0.2 and 30 possible alleles per site, except the stochastic objective-ablation collection.

| Collection | Simulation settings | Replicates | Analysis |
|---|---|---|---|
| Objective ablation (stochastic) | *n* = 500, 30 sites; *μ* = 0.36, 10 alleles; *ε* = 0, *ε*<sub>*S*</sub> = 0.25 | 25 | Objective order, quartet sampling, penalties |
| Objective ablation (mixed) | Development mixed regime, (*ε*, *ε*<sub>*S*</sub>) = (0.15, 0.10) | 20 | Objective order, quartet sampling, penalties |
| Development | *n* = 500, 30 sites; (*ε*, *ε*<sub>*S*</sub>) = (0.02, 0.20), (0.15, 0.10), (0.28, 0.02) | 3 × 20 | Initialization |
| Dimension | *n* = 250, 1,000, 5,000 at 30 sites; same missingness regimes as development | 3 × 3 × 10 (×5 at *n* = 5,000) | Embedding dimension |
| Timing | *n* = 250–10,000 at 30 sites; 15–240 sites at *n* = 1,000; *ε* = 0.02, *ε*<sub>*S*</sub> = 0.20 | 10 × 3 | Runtime and memory |

#### A11 Baseline methods and computational evaluation

We compared HERACLES with LAML v1.0.5 [11], Startle-NNI [10], VINE v0.3.5 [17], and the greedy, neighbour-joining and UPGMA solvers of Cassiopeia v2.1.0 [9]. Every method received the same character matrix, with cell rows permuted once (seed 0) to remove the ordering inherited from the simulated trees.

Default settings were used except as follows. Startle-NNI was initialized from a neighbour-joining tree and run with unweighted parsimony on four threads. VINE used seed 1 and a minimum of 2,000 iterations rather than its default 200. Cassiopeia’s neighbour joining and UPGMA used their default weighted Hamming distance.

LAML was evaluated using both its default standalone neighbour-joining initialization and the paper’s recommended pipeline, which initializes LAML from Startle-NNI run on a Cassiopeia-greedy tree. For each benchmark collection, we reported the configuration with lower mean collapsed rooted RF on datasets completed by both configurations, favoring LAML in the comparison. The standalone configuration was selected for the LAML benchmark and the recommended pipeline for the Startle and independently generated scalability benchmarks. This selection used ground-truth accuracy on the evaluation datasets.

LAML otherwise used its default EM solver, ultrametric model, uniform mutation priors and one topology search of at most 500 iterations; the pipeline used one parameter initialization, as in the authors’ benchmark. The topology search terminated after a pass without an accepted NNI move. Unlike the authors’ benchmark, we did not supply simulated mutation priors to LAML, nor any method.

Each baseline run was limited to 24 h of wall time. Runs that timed out or terminated with an error were excluded from accuracy summaries. Means were calculated over completed runs, and paired comparisons used only datasets completed by both methods (Table S4).

We evaluated computational performance using wall-clock runtime and peak resident host memory. HERACLES runtime includes distance initialization, embedding optimization and final tree reconstruction, but excludes accuracy evaluation and optional NNI. Runtime and memory scaling (Supp. Fig. S11) were measured on simulated datasets of 250–10,000 cells with 30 recording sites (three replicates per size). Each dataset was fitted twice with HERACLES, once on four CPU cores restricted to one socket and once on one NVIDIA H100 GPU, and every baseline was run on the same datasets with four CPU cores on one socket. Atlas-scale hardware is described in Section A8; PEtracer was fitted on one NVIDIA H100 GPU.

Software versions were Python 3.10, NumPy 2.2.6, SciPy 1.15.3, pandas 2.3.3, PyTorch 2.13.0, geoopt 0.5.1, DendroPy 5.0.10 and Cassiopeia 2.1.0.

#### A12 Reconstruction accuracy and statistical analysis

We assessed reconstruction accuracy using normalized rooted Robinson–Foulds (RF) distance [36]. Before evaluation, both inferred and ground-truth trees underwent mutationless-edge contraction using Cassiopeia’s ancestral-state reconstruction (Camin–Sokal parsimony, Cassiopeia v2.1.0 [9]): an edge was contracted when the child’s inferred characters equalled its parent’s, and ground-truth trees were contracted using their dropout-free character matrices when available. This avoids penalizing relationships that cannot be resolved from the available recordings; contraction was performed only for evaluation and did not modify the reconstructed phylogenies. Let *C*_inf_ and *C*_true_ denote the sets of nontrivial rooted clades of the contracted trees, excluding single cells and the full leaf set. We calculate

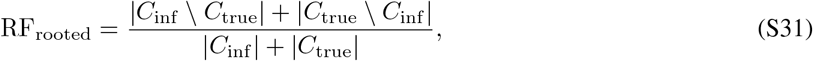

where zero indicates identical resolved clade sets and one indicates that no resolved clade is shared. The denominator counts the resolved clades of both trees rather than the number possible for *n* cells, so the normalization depends on how resolved the two trees are after contraction; if neither tree has a resolved clade, the distance is defined as zero. The same metric was used for the sequential-recording experiments.

Each method was compared separately against HERACLES within each benchmark condition. We used two-sided paired Wilcoxon signed-rank tests with Pratt’s treatment of zero differences and report unadjusted *P*-values. Tests included only datasets completed by both methods. Each significance asterisk in main Fig. 4 represents one such comparison.

The LAML benchmark included 50 paired datasets per condition, or fewer when methods failed, as indicated on the bars. The Startle benchmark included 522, 522, 346 and 346 replicates at 50, 100, 150 and 200 cells, respectively, drawn from 45, 45, 23 and 23 simulation configurations. The 1,000–10,000-cell benchmark included 60 datasets per cell count, except where methods failed to complete: the LAML pipeline completed 47 and 34 datasets, and Startle-NNI completed 47 and 49 datasets, at 1,000 and 2,000 cells, respectively.

#### A13 Analysis of KP-tracer metastatic lineages

##### Dataset and comparison trees

We analyzed tumour 3724_NT_All from the KP-tracer mouse model of metastatic lung adenocarcinoma [7]. Following LAML [11], we selected 1,461 cells with distinct edit patterns from the original 21,108 cells: 886 from the primary tumour, 449 from a soft-tissue metastasis and 9, 75 and 42 from three liver metastases. The liver metastases were numbered as in Chu et al., with site identities matched by reproducing their reported per-site migration counts.

The character matrix, tissue labels and published LAML, Startle-NNI and Cassiopeia-Hybrid trees were obtained from the LAML authors’ repository (`raphael-group/laml-experiments`, commit `9887676`). The LAML tree was reconstructed from the 1,461-cell subset, whereas the Startle-NNI and Cassiopeia-Hybrid trees were reconstructed from all 21,108 cells and pruned to the same subset. All comparison trees therefore contained identical leaves and were used without refitting.

##### HERACLES reconstruction

HERACLES was run with *D* = 192, the two-stage optimization schedule, a maximum of 5,000 steps and per-site allele probabilities, from three random seeds. Each embedding was decoded using neighbour joining, and the reconstruction with the highest fitted log-likelihood was selected without reference to tissue annotations. The three reconstructions required 107, 107 and 108 migrations after contraction.

##### Migration inference

We evaluated migration patterns on both the original trees and trees contracted at branches without inferred edits using the Camin–Sokal procedure (Section A12). Contraction removes binary resolutions unsupported by observed mutations, which can otherwise artificially reduce the minimum migration count.

For each tree, tissue labels were assigned to internal nodes by minimizing the number of transitions between sites,

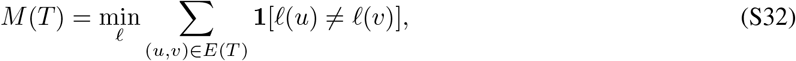

where *l* assigns tissue labels to nodes, with leaf labels fixed. We computed the minimum exactly using Sankoff small parsimony [37], with unit transition costs, as in previous analyses [10, 11].

To obtain directed migration counts, ties among optimal ancestral assignments were resolved from the root toward the leaves, favoring the primary tumour, following MACHINA [38]. Each edge connecting different tissue labels was counted as one migration from the parent’s site to the child’s. Transitions from metastatic sites back to the primary tumour were classified as reseeding events. Although the minimum total migration count is independent of the root and tie-breaking procedure, directed migration counts depend on these choices; the same procedure was applied to all trees.

#### A14 Analysis of PEtracer barcoded lineages

##### Dataset and independent lineage barcodes

We analyzed clone 4 (experiment 3) from the PEtracer barcoded-tracing experiment [3], following LAML-Pro [13]. Cells received static lentiviral barcodes on days 5 and 7, independently of the evolving lineage recorder. Each barcode marks descendants of a single ancestral integration event, providing an independent measure of lineage reconstruction accuracy.

The character matrix contains 3,108 cells and 36 recording sites arranged in 12 cassettes of three. All 13,686 missing entries occur in completely missing cassettes (4,562 cell–cassette pairs), motivating the cassette-level missing-data model (Section A9). Day-5 barcodes label 3,046 cells with 29 distinct barcodes, and day-7 barcodes label 3,018 cells with 97 distinct barcodes.

##### Integration-event metric

We evaluated each reconstruction with the LAML-Pro integration-event metric, separately for the day-5 and day-7 barcodes. Under the single-integration assumption, all carriers of a barcode should form one clade; cells in which no barcode was detected are compatible with any barcode. For each barcode *b*, let *k*_*b*_ be the number of maximal clades containing at least one carrier of *b* and no cell with a different detected barcode. The score ∑_*b*_ *k*_*b*_ counts the implied integration events, with a minimum of one per barcode (29 on day 5 and 97 on day 7). The LAML-Pro implementation computes this score by assigning barcodes to internal nodes bottom-up, propagating assignments through unlabelled nodes, and counting events on branches entering barcode-labelled clades. We verified our implementation by reproducing the published scores of the binary-refined PEtracer tree (53 and 236).

##### Comparison trees and scoring conventions

We compared HERACLES with the published PEtracer tree and a released LAML-Pro reconstruction. Because LAML-Pro requires a rooted binary starting tree, its analysis used a binary refinement of the published PEtracer tree, which contains 609 polytomies with up to 15 children. Following the LAML-Pro implementation, each polytomy with *k* children was resolved into a caterpillar with *k*− 2 additional internal nodes connected by zero-length branches, using the published child order. This retained all 1,933 original clades and added 1,174 internal nodes without additional support from the edits.

We used the highest-scoring released LAML-Pro tree, obtained from a second-stage run with error probability *η* = 0.10 and 5,000 followed by 20,000 simulated-annealing NNI moves (LAML-Pro commit `43100df`).

To match the LAML-Pro evaluation, we scored trees as distributed, including their binary resolutions. We additionally contracted branches without inferred mutations using the procedure in Section A12 and recomputed the scores. On multifurcating trees, the integration-event procedure counts separate barcoded child subtrees as distinct events, even when a binary resolution could group them. We therefore report both conventions (Table S6).

##### HERACLES reconstruction and tree selection

HERACLES was run from three random seeds using default dimension *D* = 279, per-site allele probabilities and the cassette-level missing-data model. Heritable silencing and per-cassette dropout rates were learned, both initialized at 0.001. Each embedding was decoded using neighbour joining. The reconstruction with the highest fitted log-likelihood was selected without reference to the independent lentiviral barcodes.

## B Supplementary Results

### B1 Algorithm components

#### Sensitivity to initialization

We evaluated initialization strategies on the development simulation collection (Section A10), comprising 60 datasets across three missingness regimes. HERACLES achieved similar accuracy when initialized from RapidNJ, conventional neighbour joining or MDS without a preliminary tree (Section A1; Supplementary Fig. S4). Across the three missingness regimes, differences relative to RapidNJ were at most 0.004 RF, and none was statistically significant (*P*≥ 0.29). In contrast, UPGMA initialization increased RF by 0.089, 0.079 and 0.033, and Cassiopeia-greedy initialization by 0.35, 0.26 and 0.10, in the stochastic-dominant, mixed and heritable-dominant regimes, respectively. Most fits from these poorer initializations reached the 5,000-step limit before converging, so HERACLES did not recover from substantially poorer starting trees within the default optimization budget.

**Figure S4:**
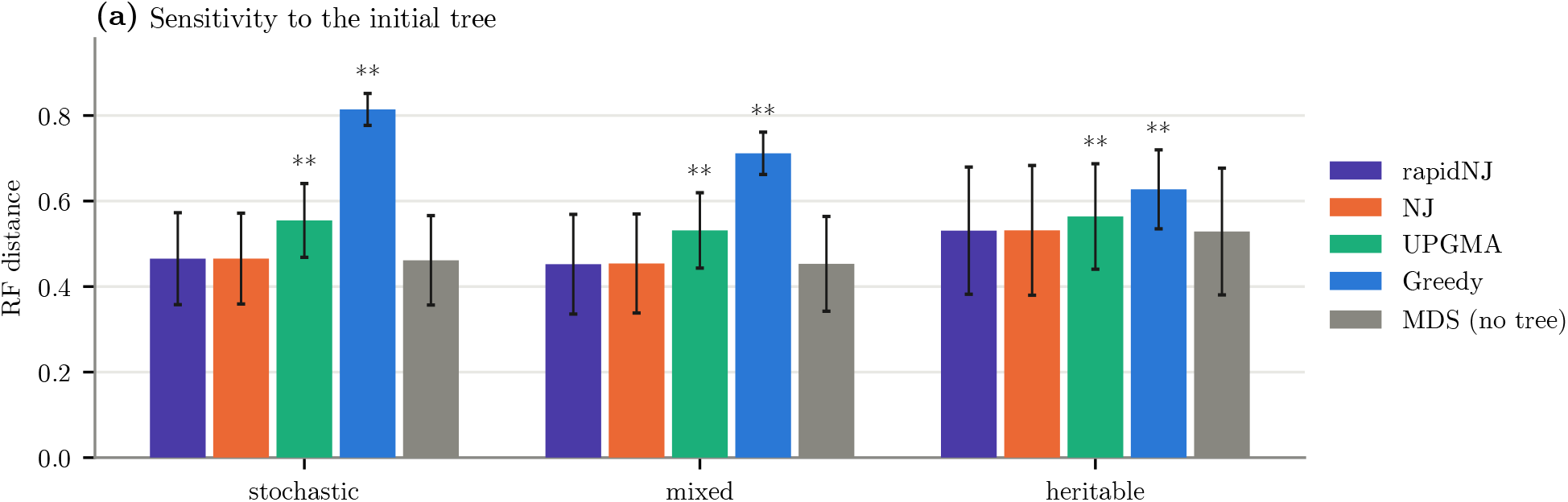
Sensitivity to initialization. Collapsed rooted RF across three missingness regimes of the development collection, using RapidNJ [33], neighbour joining [14], UPGMA [15], Cassiopeia greedy, or direct MDS initialization. Asterisks indicate two-sided paired Wilcoxon signed-rank tests against RapidNJ (^*^*P <* 0.05, ^**^*P <* 0.01).

#### Recovery of missing-data parameters

We evaluated recovery of heritable silencing and stochastic dropout rates on the LAML benchmark (Section A10; Supplementary Fig. S5). HERACLES recovered the trends in the simulated rates, although LAML estimated them more accurately (mean absolute errors of 0.022 for HERACLES and 0.009 for LAML, averaged across both missingness parameters and all 250 datasets). Estimates were similar when both rates were initialized at 0.001 or 0.1 (median absolute difference *<* 0.005), indicating limited sensitivity to starting values.

**Figure S5:**
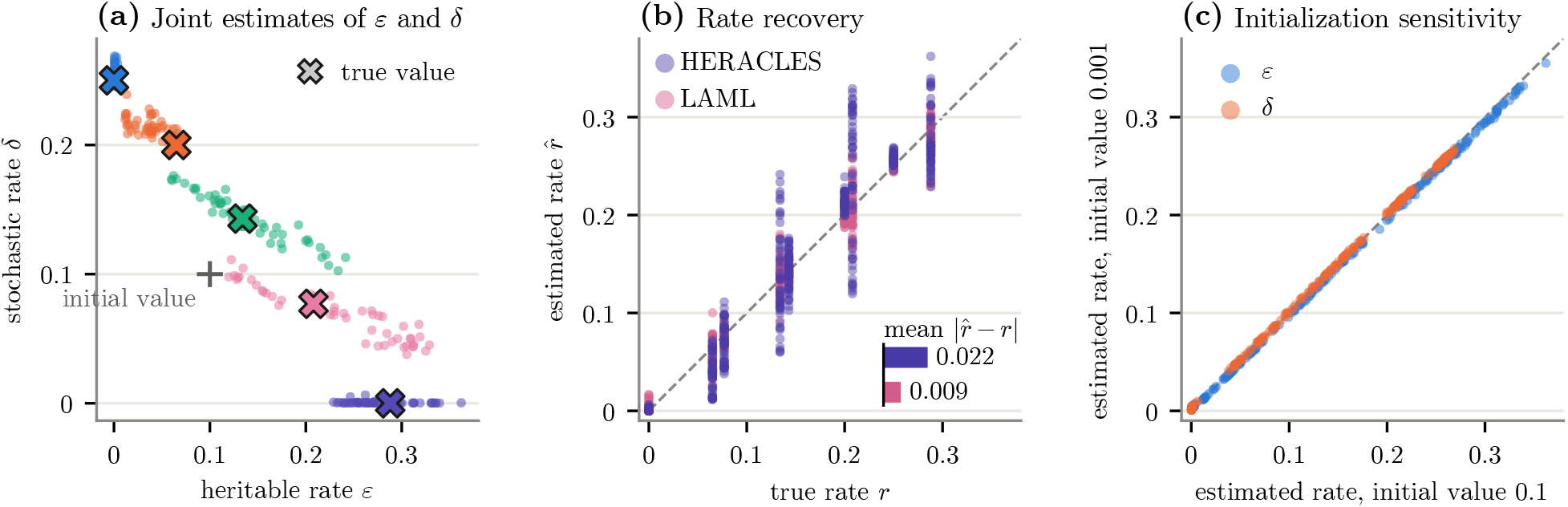
Recovery of simulated missing-data rates. **(a)** Estimated heritable (*ε*_*H*_) and stochastic (*ε*_*S*_) missingness rates on the five 250-cell conditions of the LAML benchmark (50 datasets each). **(b)** Estimated versus simulated rates for HERACLES and LAML. **(c)** HERACLES estimates from initial rates of 0.001 and 0.1.

#### Geometric regularization

We evaluated the ultrametric and four-point penalties (Section A6) on the two objective-ablation collections (Section A10; Supplementary Fig. S6), with a budget of 20,000 steps. Removing the four-point penalty increased RF by 0.018 under mixed missingness (*P* = 0.003) and by 0.003 under stochastic missingness (*P* = 0.51). Removing the ultrametric penalty increased RF by 0.019 under stochastic missingness (*P* = 2.6 × 10^−5^) and by 0.008 under mixed missingness (*P* = 0.033); without it, 3 of 25 stochastic-collection fits diverged numerically, leaving 22 matched datasets for the paired comparison. With the four-point penalty active, the mean defect measured on uniformly sampled quartets fell from 0.045–0.049 to 0.014–0.020 (Supp. Fig. S7).

**Figure S6:**
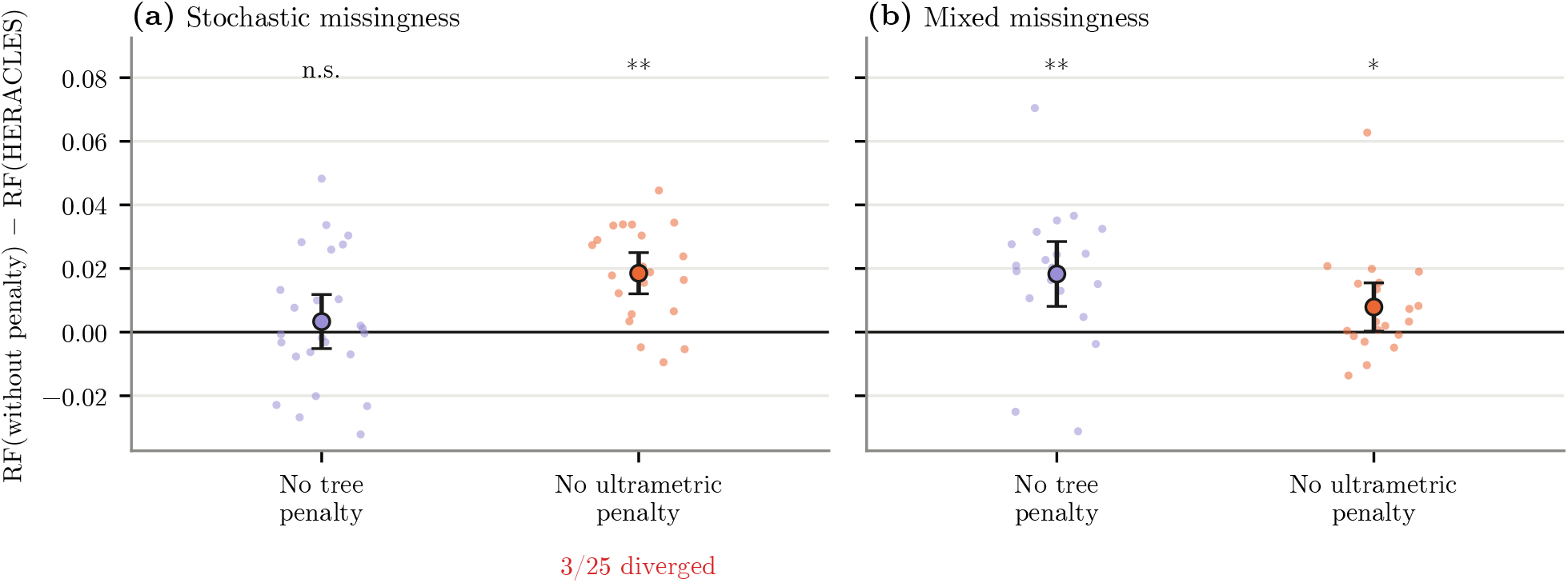
Sensitivity to the geometric penalties. Paired change in collapsed rooted RF when the four-point (“No tree penalty”) or ultrametric penalty is removed, relative to the full configuration, on the (a) stochastic (25 datasets) and (b) mixed (20 datasets) objective-ablation collections. Three stochastic-collection fits without the ultrametric penalty diverged and are excluded. Paired Wilcoxon tests: n.s., *P >* 0.05; *, *P <* 0.05; **, *P <* 0.01.

**Figure S7:**
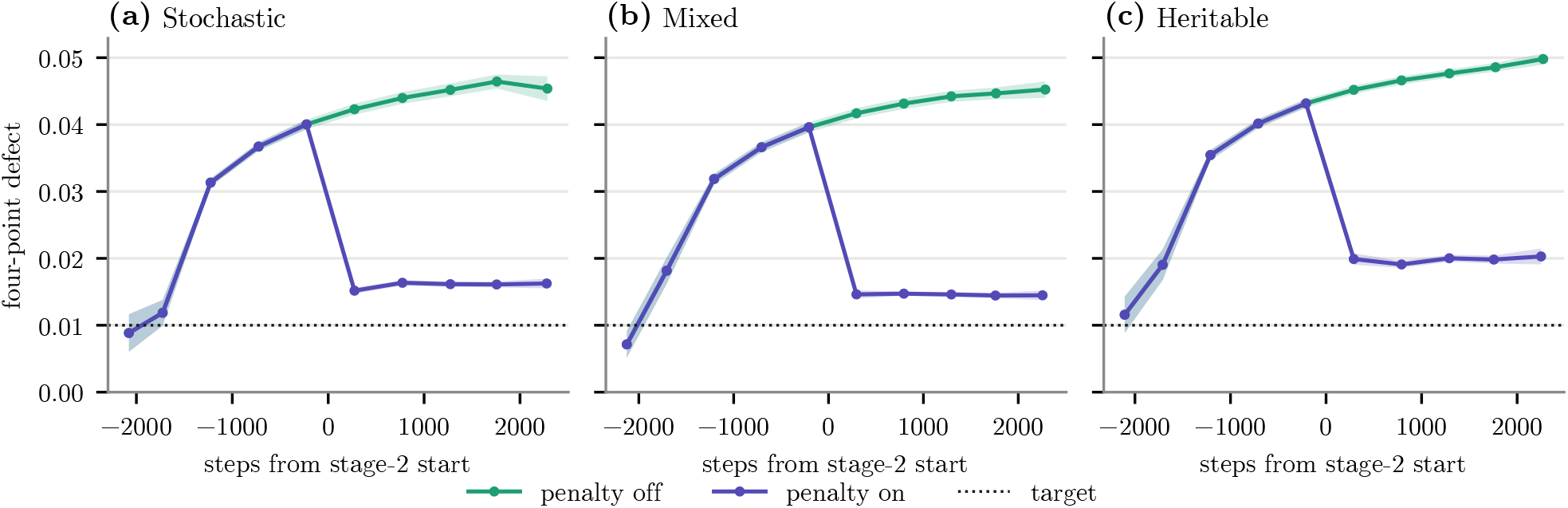
Four-point defect during optimization. Mean four-point defect on uniformly sampled quartets, aligned to the start of stage 2, for fits with and without the four-point penalty on the (a) stochastic-only (0/100), (b) mixed (25/75) and (c) heritable-only (100/0) 250-cell LAML benchmark conditions (50 datasets each).

This pattern may reflect the different phylogenetic information carried by the two missingness mechanisms: heritable silencing produces correlated missingness among descendants, whereas stochastic dropout obscures observations without providing an equivalent ancestry signal. Geometric regularization may therefore be most beneficial when the evolutionary likelihood is less strongly constrained by the observed recording data.

#### Embedding dimension

We evaluated embedding dimension using the separate dimension simulation collection (Section A10), spanning 250–5,000 cells (Supp. Fig. S8). Increasing the dimension initially improved reconstruction accuracy, but the gains diminished at larger dimensions. Improvements from doubling *D* were small and nonsignificant near the default 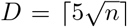 at 250 and 1,000 cells, but at 5,000 cells doubling from 256 to 512 dimensions still reduced RF by 0.0075 (*P* = 6 ×10^−5^), while optimization time increased with dimension. The default is therefore a compromise between accuracy and computational cost that becomes slightly conservative at larger *n*.

**Figure S8:**
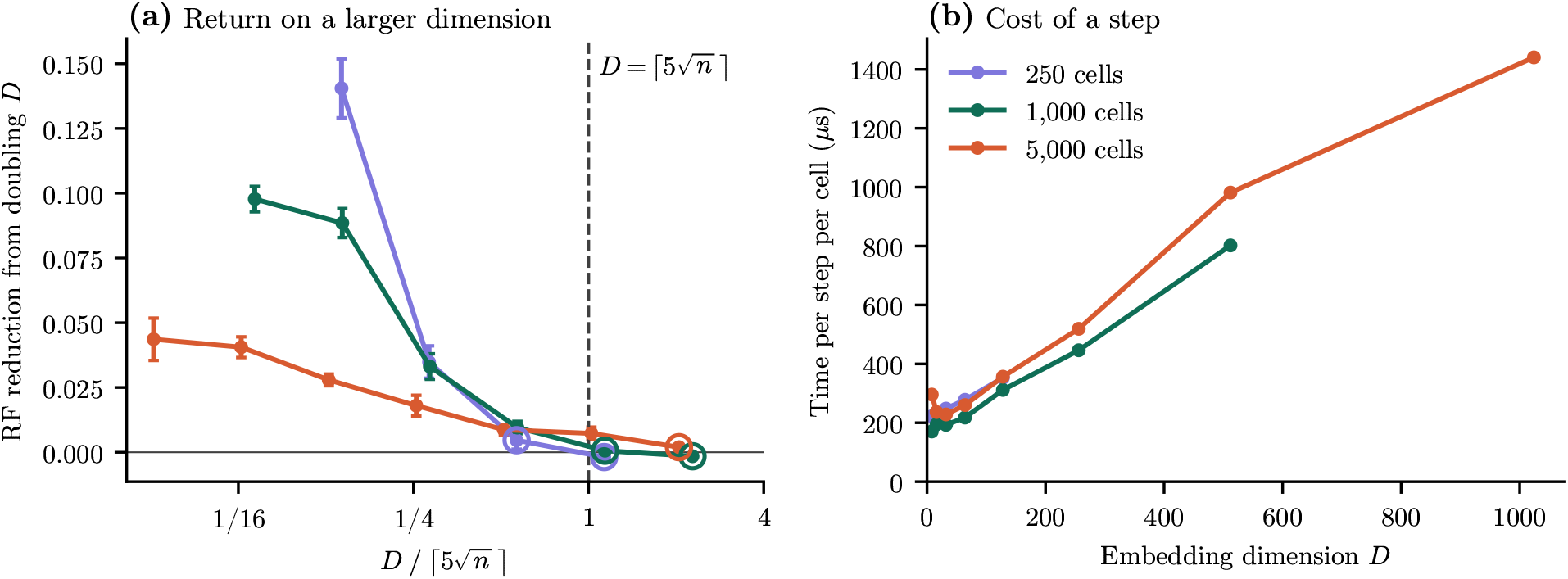
Effect of embedding dimension on accuracy and runtime. **(a)** Improvement in RF when doubling *D* (mean ±SEM), plotted relative to the default dimension 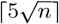. Open circles indicate nonsignificant improvements (*P >* 0.05, paired Wilcoxon test). **(b)** Optimization time per step per cell.

#### Informative sampling

A quartet carries information about a branch only when the branch splits its four cells two and two. We therefore measured, for each branch of the true tree, how many such quartets each sampler places on it per training step. Both samplers were built exactly as in a fit, from the weighted Hamming distances used for initialization, and each drew 20 batches of 15*n* quartets with three random seeds on fifteen trees of the LAML 50/50 condition (500, 750 and 1,000 cells, five each); no embedding was fitted. We grouped branches by the number of cells below them and report the mean number of informative quartets per step and the fraction of branches receiving fewer than one per step. Under uniform sampling, 93% of branches above clades of 2–3 cells and 11% above clades of 4–7 cells received fewer than one informative quartet per step. Neighbourhood-based sampling raised coverage of these branches to 13 and 25 quartets per step, leaving 3% and 0% undertested, while reducing coverage of the largest clades from 1,494 to 460 quartets per step (Supp. Fig. S9).

**Figure S9:**
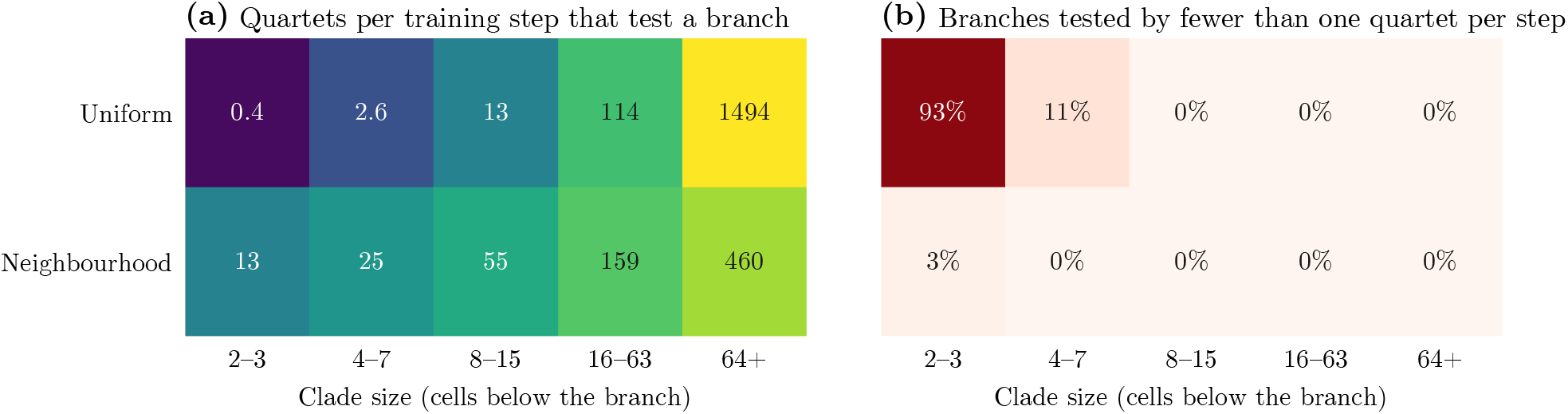
Quartet coverage by clade size. (a) Mean number of quartets per training step that test a branch, by the number of cells below the branch, for uniform and neighbourhood-based sampling. (b) Fraction of branches tested by fewer than one quartet per step. Fifteen 500–1,000-cell trees from the LAML 50/50 benchmark condition, 15*n* quartets per step.

### B2 Simulation benchmarks

#### Accuracy on published benchmarks

HERACLES achieved the lowest reconstruction error among methods without post-processing across all eight conditions of the LAML benchmark (Table S2).

LAML’s accuracy depended strongly on its initial tree. Default LAML, initialized from neighbour joining, outper-formed the authors’ recommended pipeline (Cassiopeia greedy →Startle-NNI →LAML), with mean RF of 0.345 versus 0.445 across 290 jointly completed datasets and lower error on 266 of them. On the 500–1,000-cell conditions, LAML’s topology search returned its neighbour-joining starting tree unchanged in 146 of 150 datasets. Together, these findings suggest that LAML’s discrete topology search becomes increasingly dependent on initialization as lineage size increases.

**Table S2:** LAML benchmark: mean± SD collapsed rooted Robinson–Foulds (RF) distance (lower is better) for the eight published simulation conditions. At 250 cells, setting labels give the percentage shares of heritable and stochastic missingness; the 500–1,000-cell settings use the 50/50 regime. LAML is started from the Cassiopeia NJ tree (from NJ) or from the Startle-NNI tree, as in the authors’ pipeline (from Startle). Bold marks the lowest RF in each row and italics the two lowest. Superscripts give the number of completed runs where fewer than all 50 replicates finished.

| Setting | HERACLES | LAML (from NJ) | LAML (from Startle) | Startle-NNI | VINE | NJ | UPGMA | Greedy |
|---|---|---|---|---|---|---|---|---|
| 0/100, 250 cells | ***0.357*** ± ***0.071*** | *0.377* ± *0.069* | 0.482 ± 0.090 | 0.399 ± 0.072 | 0.633 ± 0.084 | 0.447 ± 0.057 | 0.439 ± 0.066 | 0.810 ± 0.047 |
| 25/75, 250 cells | ***0.304*** ± ***0.059*** | *0.324* ± *0.067* | 0.437 ± 0.086 | 0.367 ± 0.068 | 0.515 ± 0.070 | 0.407 ± 0.059 | 0.401 ± 0.054 | 0.787 ± 0.043 |
| 50/50, 250 cells | ***0.276*** ± ***0.045*** | *0.294* ± *0.056* | 0.406 ± 0.083<sup>40</sup> | 0.358 ± 0.052<sup>40</sup> | 0.420 ± 0.066 | 0.394 ± 0.046 | 0.392 ± 0.055 | 0.749 ± 0.044 |
| 75/25, 250 cells | ***0.198*** ± ***0.051*** | *0.221* ± *0.070* | 0.313 ± 0.086<sup>20</sup> | 0.292 ± 0.054<sup>20</sup> | 0.316 ± 0.078 | 0.365 ± 0.074 | 0.374 ± 0.074 | 0.733 ± 0.052 |
| 100/0, 250 cells | ***0.117*** ± ***0.028*** | *0.152* ± *0.042* | 0.191 ± 0.094<sup>10</sup> | 0.265 ± 0.060<sup>10</sup> | 0.214 ± 0.051 | 0.327 ± 0.062 | 0.346 ± 0.057 | 0.675 ± 0.050 |
| 50/50, 500 cells | ***0.270*** ± ***0.045*** | 0.386 ± 0.040 | 0.475 ± 0.065<sup>40</sup> | *0.370* ± *0.044*<sup>40</sup> | 0.464 ± 0.047 | 0.392 ± 0.043 | 0.374 ± 0.047 | 0.741 ± 0.034 |
| 50/50, 750 cells | ***0.270*** ± ***0.040*** | 0.396 ± 0.032 | 0.493 ± 0.060<sup>40</sup> | 0.384 ± 0.033<sup>40</sup> | 0.490 ± 0.041 | 0.396 ± 0.032 | *0.370* ± *0.038* | 0.730 ± 0.024 |
| 50/50, 1000 cells | ***0.277*** ± ***0.038*** | 0.400 ± 0.028 | 0.497 ± 0.042<sup>40</sup> | 0.383 ± 0.034<sup>40</sup> | 0.509 ± 0.037 | 0.400 ± 0.028 | *0.361* ± *0.034* | 0.724 ± 0.022 |

On the Startle benchmark, HERACLES achieved the lowest error at every cell count (Table S3), with small additional improvements from NNI refinement. Its advantage was also observed across recording-site counts, editing probabilities and dropout rates (Supp. Fig. S10). Unlike on the LAML benchmark, LAML’s recommended pipeline outperformed standalone initialization (mean RF 0.350 versus 0.377 across 1,736 datasets; lower error on 1,005). We therefore used the better-performing LAML configuration for each benchmark, as described in Section A11.

**Table S3:** Startle benchmark: mean± SD collapsed rooted Robinson–Foulds (RF) distance (lower is better) by number of cells, pooled over the available published simulation configurations. LAML is started from the Cassiopeia NJ tree (from NJ) or from the Startle-NNI tree, as in the authors’ pipeline (from Startle). Bold marks the lowest RF in each row and italics the two lowest. The grid is uneven across cell counts, recording-site counts, editing probabilities and dropout rates, so these summaries are not controlled one-factor comparisons.

| Setting | HERACLES | LAML (from NJ) | LAML (from Startle) | Startle-NNI | VINE | NJ | UPGMA | Greedy |
|---|---|---|---|---|---|---|---|---|
| 50 cells | ***0.271*** ± ***0.207*** | 0.327 ± 0.249 | *0.301* ± *0.229* | 0.305 ± 0.235 | 0.439 ± 0.271 | 0.334 ± 0.246 | 0.362 ± 0.226 | 0.353 ± 0.232 |
| 100 cells | ***0.299*** ± ***0.210*** | 0.372 ± 0.247 | 0.354 ± 0.234 | *0.344* ± *0.235* | 0.518 ± 0.278 | 0.375 ± 0.242 | 0.403 ± 0.230 | 0.416 ± 0.239 |
| 150 cells | ***0.314*** ± ***0.227*** | 0.405 ± 0.272 | 0.372 ± 0.255 | *0.369* ± *0.257* | 0.549 ± 0.296 | 0.406 ± 0.270 | 0.443 ± 0.250 | 0.444 ± 0.258 |
| 200 cells | ***0.336*** ± ***0.224*** | 0.432 ± 0.276 | 0.397 ± 0.252 | *0.391* ± *0.258* | 0.577 ± 0.295 | 0.432 ± 0.272 | 0.459 ± 0.253 | 0.466 ± 0.254 |

**Figure S10:**
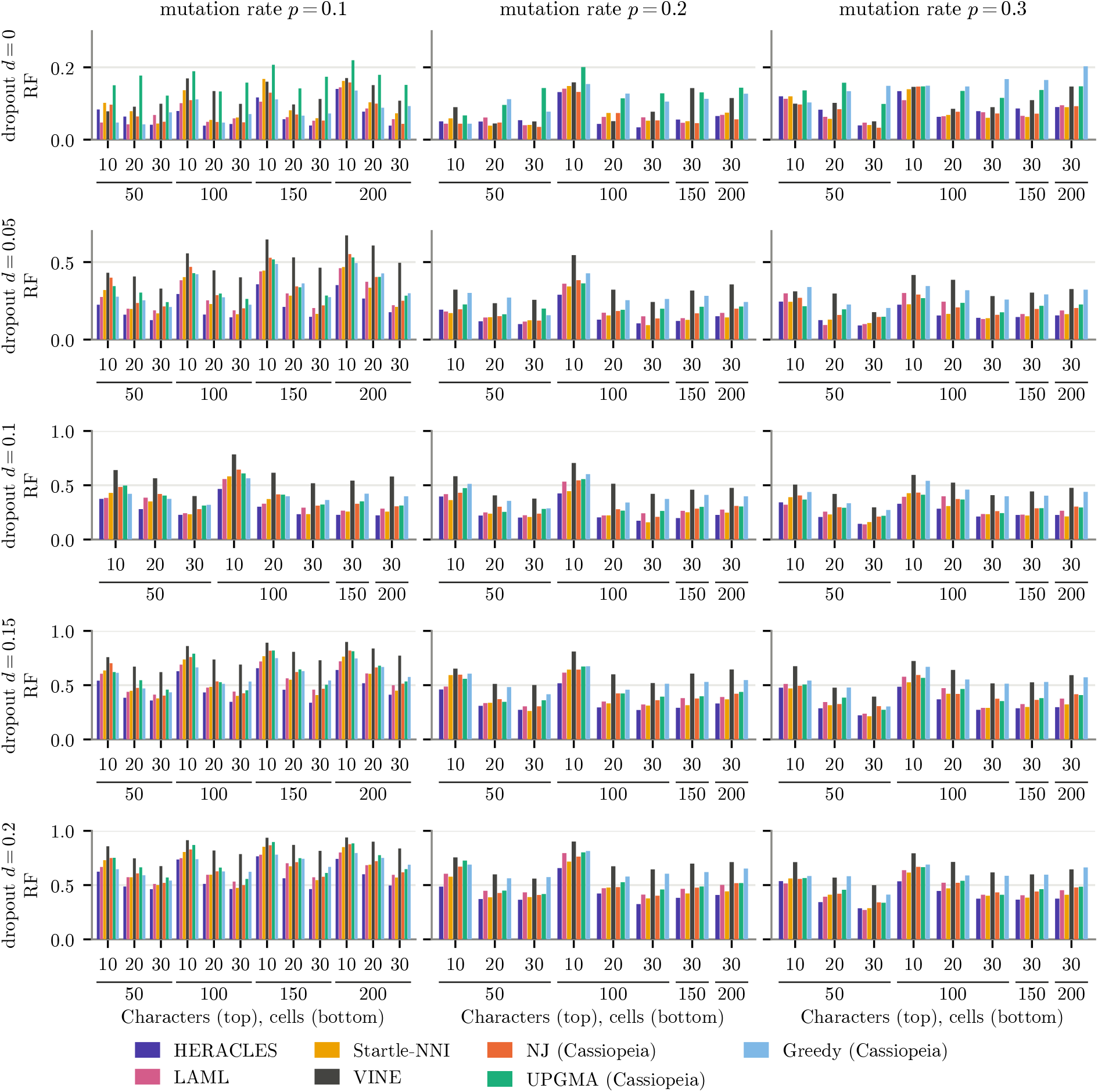
Startle benchmark accuracy by recording configuration. Collapsed rooted RF for HERACLES, LAML, Startle-NNI, VINE and Cassiopeia’s neighbour-joining, UPGMA and greedy solvers. Rows vary stochastic dropout *d* (0–0.20), columns vary the per-edge editing probability *p* (0.1–0.3), and grouped x-axis labels give the number of recording sites (10, 20 or 30) and cells (50–200). Blank combinations were not simulated in the published benchmark.

#### Accuracy at increasing lineage scales

Across the independently generated 1,000–10,000-cell simulations (Section A10), HERACLES maintained an accuracy advantage over scalable distance-based methods (main Fig. 4c). Neither LAML nor Startle-NNI completed reconstructions at 5,000 or 10,000 cells within the 24-h wall-time limit (Table S4). At atlas scale (25,000–150,000 cells), HERACLES used the low-memory implementation described in Section A8, with 8*n* quartets per step and an average-linkage read-out. It maintained its accuracy advantage over UPGMA and RapidNJ across all four lineage sizes, demonstrating that the improvements from likelihood-guided embedding optimization persist at scales beyond conventional likelihood-based phylogenetic reconstruction.

#### Computational scaling

With *O*(*n*) sampled quartets and embedding dimension 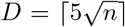, optimization cost per step is expected to scale as *O*(*n*^1.5^). The number of optimization steps was nearly constant across lineage sizes (3,600–3,733), so total runtime tracked the cost per step. On four CPU cores, total runtime scaled empirically as *n*^1.24^ across 250–10,000 cells and *n*^1.38^ across 1,000–10,000 cells (log–log fits to mean runtimes over three replicates; Supp. Fig. S11), from 3.3 min at 250 cells to 5.0 h at 10,000 cells.

**Table S4:** Benchmark completion rates. Completed reconstructions by method and condition. E indicates an error, T a 24-h timeout. LAML uses standalone initialization on its own benchmark and the recommended pipeline elsewhere.

| Panel | Condition | Replicates | HERACLES | LAML | Startle-NNI | VINE | NJ, UPGMA, greedy |
|---|---|---|---|---|---|---|---|
| LAML | 0/100, 250 cells | 50 | 50 | 50 | 50 | 50 | 50 |
|  | 25/75, 250 cells | 50 | 50 | 50 | 50 | 50 | 50 |
|  | 50/50, 250 cells | 50 | 50 | 50 | 40 (10 E) | 50 | 50 |
|  | 75/25, 250 cells | 50 | 50 | 50 | 20 (30 E) | 50 | 50 |
|  | 100/0, 250 cells | 50 | 50 | 50 | 10 (40 E) | 50 | 50 |
|  | 50/50, 500 cells | 50 | 50 | 50 | 40 (10 E) | 50 | 50 |
|  | 50/50, 750 cells | 50 | 50 | 50 | 40 (10 E) | 50 | 50 |
|  | 50/50, 1,000 cells | 50 | 50 | 50 | 40 (10 E) | 50 | 50 |
| Startle | 50–200 cells | 1,736 | 1,736 | 1,736 | 1,736 | 1,736 | 1,736 |
| Simulated | 1,000 cells | 60 | 60 | 47 (13 E) | 47 (13 E) | 60 | 60 |
|  | 2,000 cells | 60 | 60 | 34 (11 E, 15 T) | 49 (11 E) | 60 | 60 |
|  | 5,000 cells | 60 | 60 | 0 (60 T) | 0 (60 T) | 60 | 60 |
|  | 10,000 cells | 60 | 60 | 0 (60 T) | 0 (60 T) | 60 | 60 |

GPU acceleration substantially reduced runtime at larger scales: a 10,000-cell fit took 17 min on one NVIDIA H100 GPU. At atlas scale, mean GPU runtimes were 0.36, 0.97, 2.8 and 4.1 hours for 25,000, 50,000, 100,000 and 150,000 cells, respectively, compared with 8.4 and 22.5 hours on 16 CPU cores at 25,000 and 50,000 cells; two of five 50,000-cell CPU fits exceeded the 24-h limit. At atlas scale, peak host memory was dominated by the quadratic distance matrix used for final tree reconstruction rather than embedding optimization.

**Figure S11:**
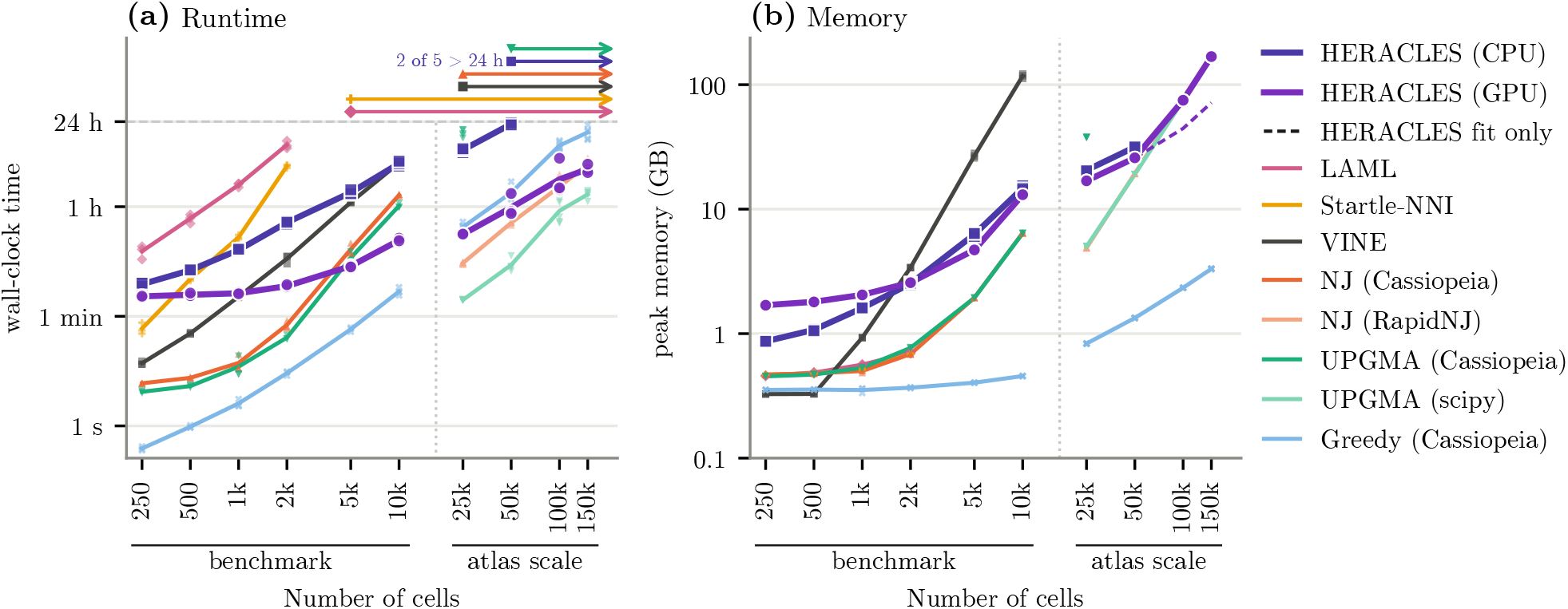
Runtime and memory scaling. **(a)** Wall-clock runtime and **(b)** peak resident host memory against cell count. The dotted line separates the timing collection (250–10,000 cells) from atlas-scale simulations (25,000–150,000 cells). HERACLES was run on four CPU cores (benchmark), 16 CPU cores (atlas scale) or one NVIDIA H100 GPU. Dots represent individual replicates; lines connect means at each cell count. Arrows indicate methods that did not complete within the 24-h runtime limit.

### B3 KP-tracer metastatic lineages

We reconstructed the KP-tracer metastatic lineage comprising 1,461 cells from five anatomical sites (Section A13). Migration estimates depended substantially on whether branches unsupported by observed edits were contracted (Table S5). Binary resolutions unsupported by observed edits impose arbitrary ancestral relationships that can artificially reduce inferred migration counts. Contracting these branches avoids attributing biological significance to unsupported topology and provides a more conservative basis for comparing migration histories.

Without contraction, LAML required the fewest migrations (99, versus 101 for HERACLES). After contraction, HERACLES required the fewest (107, versus 120 for LAML, 123 for Startle-NNI and 136 for Cassiopeia-Hybrid). HERACLES also required the fewest reseeding events under both conventions: 36 without contraction (versus 44–68) and 35 after contraction (versus 54–68).

**Table S5:** Migration counts for KP-tracer metastatic lineages. Minimum migrations inferred from tumour 3724_NT_All trees, before and after contraction of branches without inferred edits. P, primary tumour; S, soft-tissue metastasis; L1–L3, liver metastases 1–3 (9, 75 and 42 cells), numbered as in Chu et al. Totals are independent of tie-breaking; directed migration counts use ties broken toward the primary tumour. Site pairs without migrations are omitted.

|  | Tree | Total | Reseeding | P→S | S→P | P→L1 | P→L2 | L2→P | P→L3 | L3→P | S→L3 |
|---|---|---|---|---|---|---|---|---|---|---|---|
| Without contraction | Cassiopeia-Hybrid | 136 | 68 | 15 | 57 | 9 | 7 | 11 | 36 | 0 | 1 |
|  | Startle-NNI | 119 | 56 | 9 | 48 | 9 | 9 | 8 | 35 | 0 | 1 |
|  | LAML | **99** | 44 | 15 | 37 | 9 | 11 | 4 | 19 | 3 | 1 |
|  | HERACLES | 101 | **36** | 16 | 29 | 9 | 5 | 5 | 34 | 2 | 1 |
| With contraction | Cassiopeia-Hybrid | 136 | 68 | 15 | 57 | 9 | 7 | 11 | 36 | 0 | 1 |
|  | Startle-NNI | 123 | 59 | 10 | 51 | 9 | 9 | 8 | 35 | 0 | 1 |
|  | LAML | 120 | 54 | 12 | 47 | 9 | 9 | 7 | 35 | 0 | 1 |
|  | HERACLES | **107** | **35** | 18 | 30 | 9 | 5 | 5 | 39 | 0 | 1 |

Contraction particularly affected LAML, increasing its minimum migration count by 21. It also eliminated LAML’s apparent advantage for liver metastasis 3: without contraction, LAML required 22 migrations between this metastasis and the primary tumour, versus 35–36 for the other trees; after contraction, all four trees required 35–39.

### B4 PEtracer barcoded tracing

We evaluated HERACLES on 3,108 PEtracer cells using independent lentiviral barcodes introduced on days 5 and 7 (Section A14). As in the KP-tracer analysis, we contracted branches unsupported by recorder edits before evaluation, avoiding arbitrary binary resolutions that can reduce inferred integration counts. The likelihood-selected HERACLES reconstruction required 52 and 208 integrations on days 5 and 7, respectively, compared with 49 and 220 for LAML-Pro and 58 and 255 for PEtracer (Table S6). Thus, among these reconstructions, HERACLES required the fewest day-7 integrations, while LAML-Pro required the fewest on day 5.

To assess the effect of contraction, we also scored trees before contraction. Resolving the published PEtracer tree’s polytomies into arbitrary binary branches reduced its scores from 60 and 258 to 53 and 236, without adding phylogenetic evidence. Before contraction, the selected HERACLES reconstruction required 50 and 192 integrations, compared with 47 and 201 for the released LAML-Pro tree and 53 and 236 for binary-refined PEtracer. The relative ranking of these three reconstructions was unchanged by contraction. The LAML-Pro publication additionally reports scores of 42 and 195 for a final tree that was not available in the released repository and could therefore not be evaluated after contraction.

**Table S6:** Barcode integration events in PEtracer reconstructions. Inferred integration events (day 5 / day 7; lower is better) before and after contraction of branches without inferred mutations. Internal-node counts are shown under both conventions. HERACLES was fitted from three random seeds; the reconstruction shown in main Fig. 6 (†, bold) was selected by fitted evolutionary likelihood, without reference to the barcodes. The main figure reports contracted scores. The theoretical minima are 29 and 97 integration events for days 5 and 7, respectively.

| Tree | Before contraction | Contracted | Internal nodes |  |
|---|---|---|---|---|
|  |  |  | Before contraction | Contracted |
| PEtracer, binary refinement | 53 / 236 | 58 / 255 | 3,107 | 2,086 |
| PEtracer, published | 60 / 258 | – | 1,933 | – |
| LAML-Pro | 47 / 201 | 49 / 220 | 3,107 | 2,291 |
| HERACLES, seed 1 | 44 / 206 | 45 / 224 | 3,107 | 2,361 |
| **HERACLES, seed 2**<sup>†</sup> | **50 / 192** | **52 / 208** | **3,107** | **2,354** |
| HERACLES, seed 3 | 48 / 195 | 51 / 216 | 3,107 | 2,351 |

The selected HERACLES reconstruction required 6.6 min for embedding optimization on one NVIDIA H100 GPU, without NNI refinement. The released LAML-Pro reconstruction required 4.1 and 14.6 h for its two optimization stages, respectively, according to the authors’ logs from a macOS machine. These runtimes were measured on different hardware.

### B5 DNA Typewriter sequential recordings

Explicitly modeling sequential editing improved reconstruction accuracy across the 60 DNA Typewriter simulations with missing data (Section A10; Supplementary Fig. S12). Compared with HERACLES using the independent-site likelihood under otherwise identical settings, the sequential likelihood reduced mean collapsed rooted RF from 0.220 to 0.140, with lower error in 49 of 60 paired datasets (two-sided Wilcoxon signed-rank test, *P* = 4.5× 10^−8^).

Sequential HERACLES also outperformed all four distance-based methods: neighbour joining and UPGMA using cassette-aware distances (mean RF 0.197 and 0.224), and the original Typewriter neighbour-joining and UPGMA methods (0.341 and 0.462), with lower error in all 60 datasets for each. All four comparisons were significant (*P <* 10^−10^, two-sided paired Wilcoxon tests across all 60 datasets).

**Figure S12:**
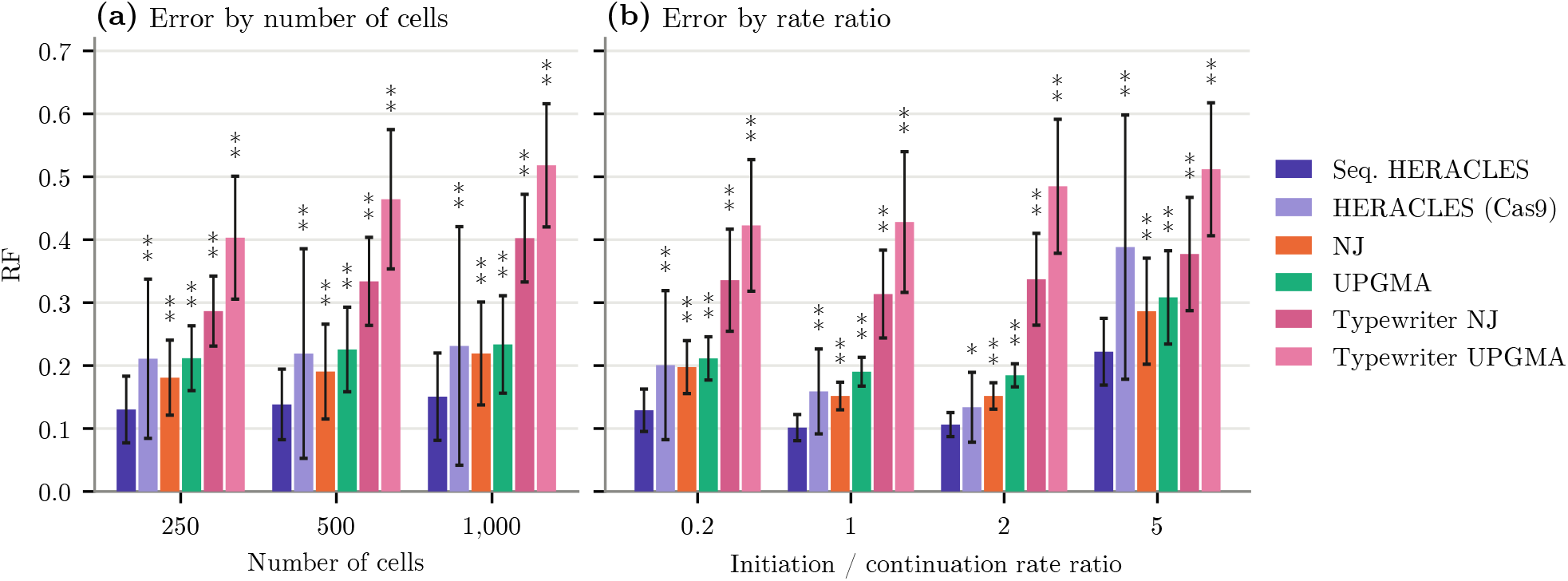
Reconstruction accuracy with a sequential DNA Typewriter recording model. **(a)** Accuracy by cell count and **(b)** by initiation-to-continuation rate ratio. Bars show mean ± standard deviation (20 and 15 datasets per setting, respectively). Asterisks indicate two-sided paired Wilcoxon signed-rank tests against sequential HERACLES within each plotted group (^*^*P <* 0.05, ^**^*P <* 0.01; unadjusted).

